# Genome wide-association study identifies novel loci in the Primary Open-Angle African American Glaucoma Genetics (POAAGG) study

**DOI:** 10.1101/2020.02.27.968156

**Authors:** Harini V. Gudiseva, Shefali Setia Verma, Venkata R. M. Chavali, Rebecca J. Salowe, Anastasia Lucas, David W. Collins, Sonika Rathi, Jie He, Roy Lee, Sayaka Merriam, Anita S. Bowman, Caitlin P. McHugh, Michael C. Zody, Maxwell Pistilli, Naira Khachataryan, Ebenezer Daniel, Windell Murphy, Mark Weiner, Jeffrey Henderer, Ahmara Ross, Qi N. Cui, Victoria Addis, Amanda Lehman, Eydie Miller-Ellis, Prithvi S. Sankar, Rohit Varma, Scott M. Williams, Gui-shuang Ying, Jason H. Moore, Marylyn D. Ritchie, Joan M. O’Brien

## Abstract

Primary open-angle glaucoma (POAG), the leading cause of irreversible blindness worldwide, disproportionately affects African Americans. Large-scale POAG genetic studies have focused on individuals of European and Asian ancestry, limiting our understanding of disease biology. Here we report genetic analysis of the largest-ever deeply phenotyped African American population (n=5950), identifying a novel POAG-associated SNP on chromosome 11 near the *TRIM66* gene (rs112369934). POAG trait association also implicated SNPs in genes involved in trabecular meshwork homeostasis and retinal ganglion cell maintenance. These new loci deepen our understanding of the pathophysiology of POAG in African Americans.

## Introduction

Primary open-angle glaucoma (POAG) is an insidious neurodegenerative disease of the optic nerve that causes progressive loss of vision.^1^ This disease affects 44 million individuals worldwide, with a projected prevalence of 53 million cases by 2020 and 80 million cases by 2040.^2^ It is estimated that up to 6 million individuals will be bilaterally blind from POAG by 2020.^3^ A disproportionately large percentage of these individuals will be of African descent.^3^ African Americans are four to five times more likely be affected by POAG than European Americans^4^—and up to 15 times more likely to experience vision loss from the disease.^5^ This population is also diagnosed at a younger age with more severe and rapidly progressive symptoms.^6,7^

Aside from African ancestry, the major risk factors for POAG include advanced age, elevated intraocular pressure (IOP), and family history of glaucoma.^8^ Several studies have also reported greater disease risk among males.^9–11^ Of these risk factors, elevated IOP is the only treatable component of the disease, with current treatments aiming to lower IOP to a level that slows or halts optic nerve degeneration.^12,13^ However, this therapeutic approach is frequently not sufficient to halt disease progression, suggesting that POAG has additional underlying disease mechanisms that could be elucidated by large-scale genetic studies.^14,15^

Many twin and family history studies have established that POAG has a strong genetic component,^16–19^ yet more than 90% of its genetics remain unexplained.^20^ This remaining 10% is due to variants with small individual effect sizes in genes such as *MYOC, OPTN, WDR36, CDK2N2B-AS1, TMC01, SIX6, ABCA1, GAS7, FOXC1, NTF4, ASB10, EFEMP1,* and *IL20RB.^20–23^* However, most of these loci do not replicate in African Americans. The majority were discovered in populations of European,^24–28^ Chinese^29^, and Japanese^30–34^ descent or large meta-analyses of these groups,^23,28,35^ and have greatly reduced or unknown role in African Americans.^36–38^

A limited number of studies have investigated glaucoma genetics in African Americans. A multi-ethnic genome-wide association study (GWAS) of the Genetic Epidemiology Research in Adult Health and Aging (GERA) cohort, which included individuals of African, Asian, Hispanic, and European descent, identified three novel and replicated variants near the *PDE7B, TMCTC2,* and *FMNL2* genes.^39^ A GWAS of South African and African American populations identified a novel candidate locus on the *EXOC4* gene, but it did not associate in a West African replication cohort.^37^ The African Descent and Glaucoma Evaluation Study (ADAGES) identified a novel association with advanced POAG and the *EN04* locus.^40^ Most recently, a GWAS identified a novel locus (*APBB2*) that was significantly associated with POAG in individuals of African ancestry, but not of European or Asian ancestry.^41^ These studies have made important strides towards understanding POAG genetics in African ancestry individuals. They do not, however, address the need to investigate the disease in a deeply phenotyped cohort of African Americans.

Since POAG is a heterogeneous disease with a broad range of trait expression, it is also important to identify variants associated with specific endophenotypes. These findings can help to identify subgroups of disease and improve predictive models for development and progression.^42,43^ To date, several associations with structural features of the optic nerve head (such as cup-to-disc ratio, rim area, cup area, and total disc area) have been reported in genes such as *CDKN2B, SIX1/6, ATOH7, CDC7,* and *CHEK2*.^44–47^ IOP has also been linked to a number of SNPs, particularly near the *TMCO1, CAV1, CAV2, FNDC3B,* and *GAS7* genes.^48–50^ Finally, central corneal thickness (CCT) has shown associations with *FOX01, ZNF469, COL5A,* and *FNDC3B*.^51–53^ As in the genetic studies on POAG described above, the variants associated with quantitative traits were primarily identified in populations of European ancestry and require explicit investigation in African Americans.

We conducted a large-scale GWAS to identify variants of pathophysiological importance to POAG in African Americans (n=5950) recruited from the city of Philadelphia, through a project entitled the Primary Open-Angle African American Glaucoma Genetics (POAAGG) study.^54^ This paper reports novel GWAS findings from overall case-control analysis, as well as genes associated with POAG endophenotypes. In addition, the functional relevance of significant findings from our study are verified using *in silico* analysis and *in vitro* studies in cell lines relevant to POAG.

## Results

### Case-control GWAS identifies potential novel associations with POAG

We successfully genotyped 6525 DNA samples on the Multi-Ethnic Genotyping Array (MEGA) V2 (EX) consortium chip and Infinium iSelect platform using Illumina FastTrack Services (Illumina, San Diego, CA) (Methods). Genotyped samples and variants were subjected to rigorous quality control (QC), as illustrated in Extended Data Tables 1 and 2. Single SNP association analysis was performed on 5950 unrelated individuals, consisting of 2589 cases (males n=1103; females n=1486) and 3361 controls (males n=1042; females n=2319). Through a logistic regression model adjusted for age, sex, and population stratification, we identified a novel locus that reached genome-wide significance in all subjects on Chromosome (Chr.) 11:8702073 [Chr. 11p15.4, rs112369934; *P* value = 2.01 × 10^-8^; adjusted odds ratio (OR)=1.67], located nearest to the *TRIM66* gene (Figure 1a).

**Fig. 1:**
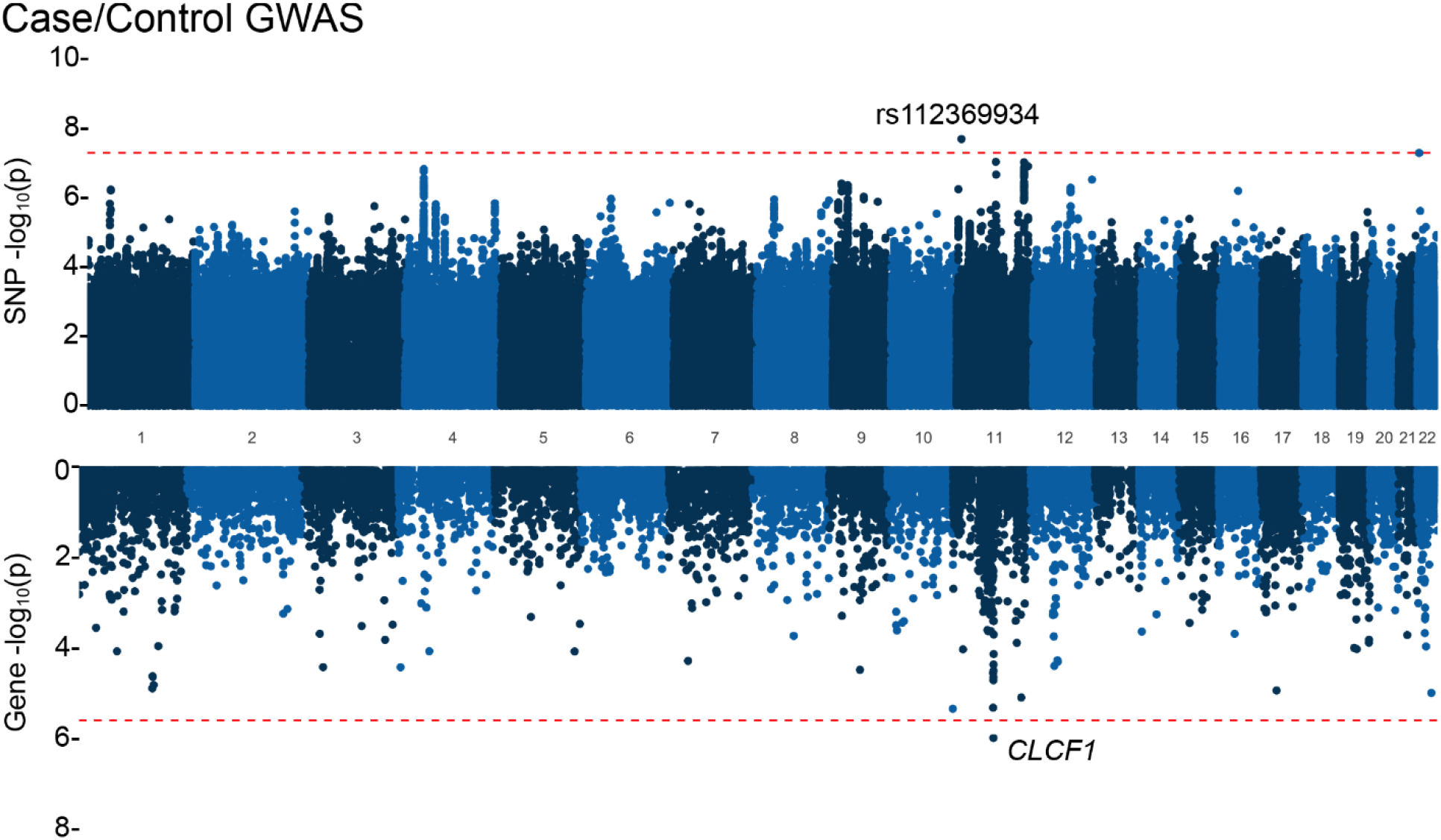
Genome-wide manhattan plots for case-control analyses. These analyses include results from case control analysis. The SNP association results are displayed in the top portion of each plot, while the association results from gene-based analyses using MAGMA are shown in the bottom portion. Genomewide Manhattan plots display –log_10_(p-values) of the logistic regression model testing the association of glaucoma status, stratified by sex, in the set of imputed variants with MAF>1%. The dotted line indicates genome-wide significance, considered here as 5×10^-8^. Gene-based Manhattan plots display –log_10_(p-values) from MAGMA analyses where SNPs were mapped to 18,991 protein coding genes. The dotted line represents Bonferroni corrected p-value threshold (p=0.05/18991=2.63e-06).

We also investigated the association of variants in high linkage disequilibrium (LD) within the gene boundaries of previously associated GWAS SNPs for POAG^28,46,55–57^ (Fig. 2). Minimal significance was observed for *CDKN2B-AS1* (lowest *P*-value=4.31 × 10^-7^; OR = 0.65) and with the downstream region of *ARHGEF12* (lowest *P*-value=9.4 × 10^-8^; OR=1.27) (Fig. 2).

**Fig. 2:**
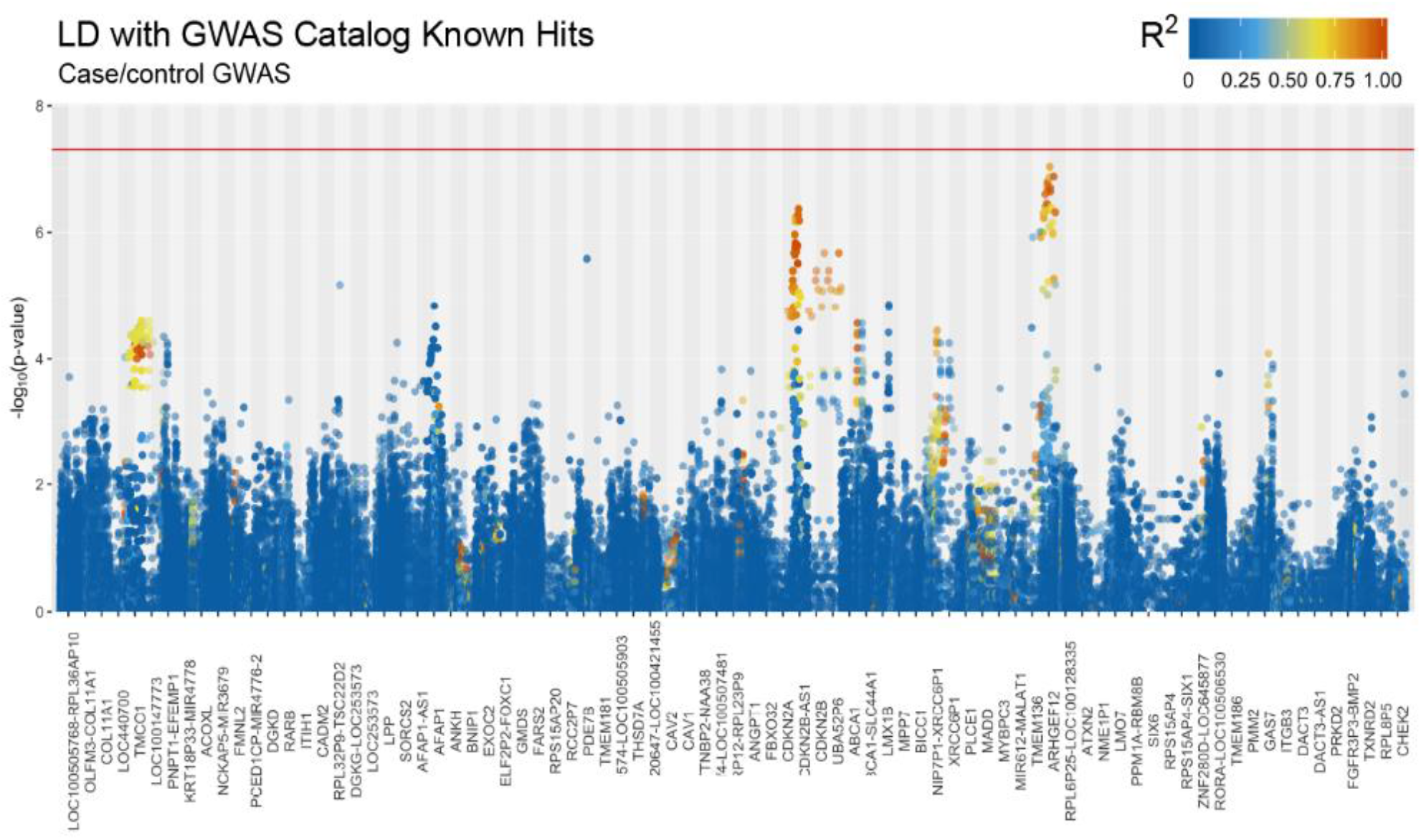
Manhattan plots displaying –log10 (p-value) of known POAG-associated genes. The points represent –log10 of p-values of known GWAS hits in case-control POAG analyses.The gradient of each colored point refers to the pairwise linkage disequilibrium R2 with GWAS reported loci.

### Quantitative traits highlight genetic heterogeneity of POAG in African Americans

Because POAG is a heterogeneous disease with diverse clinical presentations, we also conducted association analyses with glaucoma-related quantitative phenotypes for both eyes in cases and controls (Methods). Assuming independence of tests, the genome-wide quantitative trait association studies of seven distinct POAG phenotypic features identified 260 SNP-trait associations (p-value<5×10^-8^) in 127 genomic regions (Extended Table 3). A total of 30 variants reached genome-wide significance for cup-to-disc ratio (CDR), as well as 21 variants for intraocular pressure (IOP) baseline, 30 for IOP maximum (the highest IOP measure available in baseline and latest data), 43 variants for baseline pattern standard deviation (PSD), 129 variants for baseline mean deviation (MD), and one variant for retinal nerve fiber layer (RNFL) thickness (Fig. 3). SNP association results for visual acuity (VA logMAR case-control association analysis) and central corneal thickness (CCT) did not reach conservative genome-wide wide significance, assuming independence of variants and all phenotypes evaluated in this study.

**Fig. 3:**
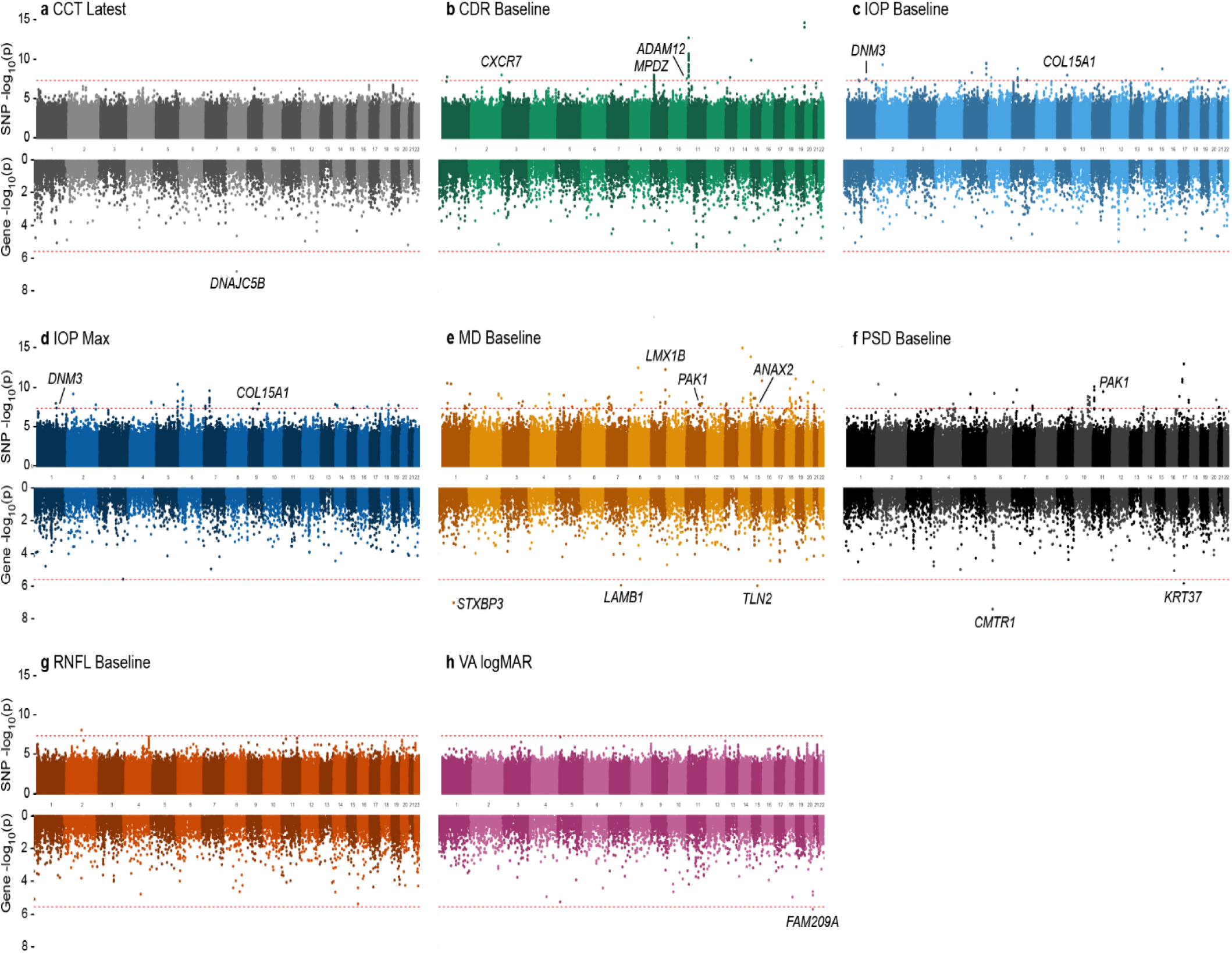
Genome-wide manhattan plots for quantitative trait analyses. The plots are color-coded by each quantitative trait, including **(a)** central corneal thickness (CCT) latest, **(b)** cup-to-disc ratio (CDR) baseline, **(c)** IOP (intraocular pressure) baseline, **(d)** IOP maximum, **(e)** mean deviation (MD) baseline, **(f)** pattern standard deviation (PSD) baseline, **(g)** retinal nerve fiber layer (RNFL) baseline, and **(h)** VA (visual acuity) in LogMAR. The SNP association results are displayed in the top portion of each plot, while the association results from gene-based analyses using MAGMA are shown in the bottom portion. Genome-wide Manhattan plots display –log_10_ (p-values) of the GEE model testing the association of the traits with the set of imputed variants with MAF>1%. The dotted line indicates genome-wide significance, considered here as 5×10^-8^. Gene-based Manhattan plots display –log_10_ (p-values) from MAGMA analyses where SNPs were mapped to 18,991 protein coding genes. The dotted line represents Bonferroni corrected p-value threshold (p=0.05/18991=2.63e-06).

A subset of the significant SNPs (n=18) mapped to genes with published POAG and phenotype association and localization in ocular tissue data, as highlighted in Table 1. We identified several genes associated with the CDR phenotype at *CXCR7/ACKR3*, *MPDZ*, and *ADAM12*. IOP baseline and maximum were both associated with *COL15A1* and *DNM3*. We also identified significant SNPs associated with MD in the *LMX1B* gene and *COL19A1* gene. Two different variants in the *PAK1* gene were independently associated with the MD and PSD phenotypes (Table 1, Extended Table 3).

**Table 1.** Selected SNPs/genes of interest associated with quantitative traits.

| Phenotype | Chr.Pos | mapped/<br>nearest<br>gene* | rsID | p-value | Beta | 95%CI | MAF |
| --- | --- | --- | --- | --- | --- | --- | --- |
| CDR | 11:3521989_T | *RPS3AP39* | rs7110375 | 1.83E-13 | -0.06 | (-0.08, -0.05) | 0.068 |
|  | 2:237653539_G | *CXCR7* | rs12328841 | 1.05E-08 | -0.03 | (-0.03, -0.02) | 0.422 |
|  | 9:13173885_G | *MPDZ* | rs4740546 | 1.06E-08 | 0.11 | (0.07, 0.15) | 0.014 |
|  | 9:13194629_T | *MPDZ* | rs1831533 | 1.63E-08 | 0.11 | (0.07, 0.15) | 0.013 |
|  | 11:3567134_A | *RPS3AP39* | rs146887003 | 1.70E-08 | -0.08 | (-0.1, -0.05) | 0.027 |
|  | 9:13181407_A | *MPDZ* | rs9696052 | 2.22E-08 | 0.11 | (0.07, 0.14) | 0.015 |
|  | 10:127738557_G | *ADAM12* | - | 2.89E-08 | 0.03 | (0.02, 0.03) | 0.411 |
|  | 9:13183256_A | *MPDZ* | rs10733255 | 3.18E-08 | 0.11 | (0.07, 0.15) | 0.014 |
| IOP<br>atBaseline<br>(mmHg) | 9:101816103_A | *COL15A1* | rs116365511 | 1.11E-08 | -1.47 | (-1.98, -0.97) | 0.011 |
|  | 9:101815916_A | *COL15A1* | rs114377702 | 1.11E-08 | -1.47 | (-1.98, -0.97) | 0.011 |
|  | 1:172221650_A | *DNM3* | rs140007960 | 3.35E-08 | -1.14 | (-1.54, -0.73) | 0.025 |
| Maximum<br>IOP (mmHg) | 9:101816103_A | *COL15A1* | rs116365511 | 1.21E-08 | -1.43 | (-1.93, -0.94) | 0.011 |
|  | 9:101815916_A | *COL15A1* | rs114377702 | 1.21E-08 | -1.43 | (-1.93, -0.94) | 0.011 |
|  | 1:172221650_A | *DNM3* | rs140007960 | 4.19E-08 | -1.17 | (-1.59, -0.75) | 0.025 |
| MD | 9:129449650_C | *LMX1B* | rs187699205 | 5.28E-13 | 3.51 | (2.56, 4.47) | 0.012 |
|  | 6:70670418_C | *COL19A1* | rs115364550 | 3.76E-08 | 3.75 | (2.41, 5.08) | 0.011 |
|  | 11:77190251_C | *PAK1* | rs145707826 | 1.62E-08 | 3.82 | (2.49, 5.15) | 0.013 |
| PSD | 11:77190646_G | *PAK1* | rs370650303 | 3.48E-10 | -1.44 | (-1.89, -0.99) | 0.013 |

\*Mapped/nearest gene=variant in gene region, with nearest gene shaded
rsID: dbSNP ID, Beta=effect size, CI=confidence interval, CDR=cup-to-disc ratio, IOP=intraocular pressure, MD=mean deviation, PSD=pattern standard deviation.

### Gene-set association analysis using MAGMA

We performed gene-based analysis using the MAGMA tool to identify additional regions of interest. Gene-based p-values were calculated based on mean SNP-association p-values. This analysis indicated *CLCF1* (p-value=1.04 x 10^-6^) as a significant gene for POAG after multiple hypothesis correction (Figure 1a). Quantitative traits-based MAGMA analyses identified six significant associations among all traits (Figure 3). CCT latest was associated with *DNAJC5B* (gene-based *P*-value=1.61 x 10^-7^) and VA logMAR case control status was associated with *FAM209A* (gene-based *P*-value=1.88 x 10^-6^). *STXBP3* (gene-based *P*-value=9.05 x 10^-8^)*, LAMB1* (gene-based *P*-value=1.07 x 10^-6^), and *TLN2* (gene-based P-value=1.0 x 10^-6^) were associated with baseline MD, and *CMTR* (gene-based *P*-value=3.74 x 10^-8^) and *KRT37* (gene-based *P*-value=1.46 x 10^-6^) were associated with baseline PSD. None of the gene-set based association results overlapped with the genes mapping to SNPs from the single variant association tests.

### In silico analysis suggests genes with possible role in POAG

To understand and prioritize POAG-associated SNPs and genes for additional studies, we assessed the expression levels of genes containing significantly associated variants *in silico* wherever data were available. We used two publicly available databases: 1) Ocular Tissue Database (OTDB) with human eye tissue expression,^58^ and 2) the IOP-induced mouse optic nerve head (ONH) microarray gene expression dataset [Glaucoma Discovery Platform (GDP)].^59^ As there is no ocular tissue information in GTEx or ENCODE databases, we did not utilize them for determining gene expression. In the OTBD, all SNP-associated genes from our study were expressed in eye tissues, including trabecular meshwork (TM) cells, the ONH, optic nerve (ON) tissues, and the retina (Table 2a). Compared to other genes, *MPDZ* had the highest expression in the majority of ocular tissues, specifically in the lens, ON, ONH, and TM, while *PAK1* expression was lowest in the ON, ONH, and TM. *COL15A1, DNM3,* and *LMX1B* were moderately expressed in all ocular tissues.

**Table 2.** Assessment of the expression, localization, and differential expression of GWAS-associated SNPs/genes using *in silico* analysis. **Table 2a:** Expression of genes in each locus that comprise over 95% of GWAS-associated gene variants in adult human eye tissues, as obtained from the Ocular Tissue Database (OTDB), along with gene expression values. In the OTDB, gene expression is represented as an Affymetrix Probe Logarithmic Intensity Error (PLIER) number for an individual probe. These numbers were calculated as described in Wagner *et al*., 2013.

| Gene | Probe ID | Choroid RPE | Ciliary Body | Cornea | Iris | Lens | Optic Nerve | Optic Nerve Head | Retina | Sclera | Trabecular Meshwork |
| --- | --- | --- | --- | --- | --- | --- | --- | --- | --- | --- | --- |
| **COL15A1** | 3.00E+06 | 34.55 | 27.33 | 35.93 | 27.9 | 32.12 | 32.44 | 32.54 | 39.29 | 35.64 | 31.97 |
| **DNM3** | 2.00E+06 | 11.68 | 5.97 | 14.67 | 14.04 | 19.67 | 25.98 | 22.12 | 30 | 17.13 | 25.1 |
| **ADAM12** | 3.00E+06 | 23.5 | 22.77 | 25.86 | 23.35 | 19.28 | 26.57 | 20.95 | 23.33 | 24.8 | 34.98 |
| **MPDZ** | 3.00E+06 | 85.28 | 64.35 | 26.63 | 58.37 | 122.56 | 165.49 | 166.36 | 93.26 | 86.54 | 121.89 |
| **LMX1B** | 3.00E+06 | 29.66 | 25.72 | 40.25 | 23.48 | 24.9 | 27.8 | 26.9 | 27.43 | 32.41 | 30.61 |
| **PAK1** | 3.00E+06 | 25.15 | 19.43 | 12.99 | 40.47 | 22.39 | 19 | 17.48 | 25.93 | 17.15 | 19.69 |

Using GDP, we compared the transcriptomic profiles of the mouse ONH obtained from 9-month-old D2 mice (an age-dependent model of IOP/glaucoma) with D2-Gpnmb+ mouse (control mice). We found increased expression of *Adam12, Cxcr7, Col15a1,* and *Anax2* in all stages of IOP-induced glaucoma, with almost two-fold overexpression in late stages (Table 2b). The expression of gene transcripts for *Trim66, Dnm3, Pak1, Mpdz,* and *Col19a1* was lower when compared to the control mice in the early to moderate stages of glaucoma, while higher in stage 5 of glaucoma (Table 2b). The differential expression patterns of the above phenotype-associated genes during moderate to advanced stages of glaucoma indicates an important role in glaucoma pathogenesis.

**Table 2b:** Gene expression levels of genome-wide associated variants in mouse ONH

| Gene | Probe Set ID | Probe Sets# | STAGE 1 |  | STAGE 2 |  | STAGE 3 |  | STAGE 4 |  | STAGE 5 |  |
| --- | --- | --- | --- | --- | --- | --- | --- | --- | --- | --- | --- | --- |
|  |  |  | FC | QV | FC | QV | FC | QV | FC | QV | FC | QV |
| *Trim66* | 1456967_at | 1 | -1.040 | 0.758 | -1.040 | 0.453 | -<br>1.354 | 0.026 | -1.596 | 0.008 | -1.370 | 0.015 |
| *Dnm3* | 1436624_at | 5 | -1.286 | 0.183 | -1.624 | 0.002 | -1.67 | 0.002 | -2.887 | 0.000 | -2.544 | 0.000 |
| *Adam12* | 1421171_at | 2 | 1.497 | 0.134 | 1.890 | 0.002 | 1.513 | 0.082 | 2.725 | 0.001 | 2.159 | 0.002 |
| *Pak1* | 1450070_s_at | 3 | -1.124 | 0.234 | -1.321 | 0.008 | -<br>1.359 | 0.011 | -1.950 | 0.001 | -1.983 | 0.000 |
| *Cxcr7* | 1417625_s_at | 1 | 1.242 | 0.409 | 1.344 | 0.036 | 1.343 | 0.062 | 2.025 | 0.003 | 1.660 | 0.008 |
| *Rbfox1* | 1418314_a_at | 1 | -1.139 | 0.520 | -1.465 | 0.005 | -<br>1.589 | 0.011 | -2.405 | 0.002 | -3.741 | 0.000 |
| *Col15a1* | 1448755_at | 1 | 1.182 | 0.548 | 1.798 | 0.002 | 1.408 | 0.027 | 2.041 | 0.004 | 1.914 | 0.000 |
| *Col19a1* | 1421698_a_at | 3 | -1.501 | 0.296 | -1.506 | 0.037 | -<br>1.128 | 0.372 | -1.947 | 0.013 | 1.090 | 0.369 |
| *Macrod2* | 1432431_s_at | 4 | -1.061 | 0.729 | 1.062 | 0.397 | 1.64 | 0.001 | 1.302 | 0.015 | 1.246 | 0.062 |
| *Mpdz* | 1418664_at | 4 | -1.057 | 0.627 | -1.124 | 0.074 | -1.14 | 0.060 | -1.091 | 0.232 | 1.070 | 0.281 |
| *Lmx1b* | 1421769_at | 1 | -1.100 | 0.540 | 1.013 | 0.518 | -<br>1.082 | 0.227 | -1.003 | 0.562 | -1.042 | 0.379 |
| *Anxa2* | 1419091_a_at | 1 | 1.332 | 0.029 | 1.625 | 0.000 | 1.969 | 0.000 | 2.051 | 0.000 | 1.830 | 0.000 |

The above table shows previously published microarray gene expression dataset in optic nerve head punches and was obtained by submitting data into the OTDB database server. The Log2 fold change (FC) shown here compares DBA/2J mice relative to DBA/2J-Gpnmb<sup>+</sup> controls at 5 molecularly distinct stages of glaucoma defined using hierarchical clustering. Stages 1, 2 and 3 are early states of glaucoma, and Stages 4 and 5 have eyes with moderate and severe optic nerve damage, respectively. Q-value (QV) shows the significance of the FC of GWAS genes at each stage.

Among the genes identified by MAGMA analysis, higher expression of *STXBP3* was observed in all ocular tissues compared to other genes, specifically in the lens, ON, ONH, and TM (Extended Table 4a). *LAMB1* was predominantly expressed in TM and cornea tissue. Differential upregulation of *Clcf1* was seen in Stage 3 and Stage 4 glaucoma in the mouse ONH. *Tln2* (FDR, q<0.05) was found to be differentially downregulated in glaucoma from early to severe stages of the disease in the mouse ONH. None of the other MAGMA identified genes associated with POAG endophenotypes showed differential expression across any stage of glaucoma (Extended Data Table 4b).

### Expression of POAG-associated loci/genes in ocular cell lines after oxidative stress

The pathogenesis of POAG involves apoptosis of retinal ganglion cells (RGCs) and damage to the TM caused by oxidative stress.^60,61^ To understand expression patterns and determine the functional relevance of POAG-associated genes under oxidative stress conditions, we induced oxidative stress in human ocular cell lines with H202 (Table 3). We quantified the expression profiles of POAG-associated genes in human TM cells (hTM) and RGCs derived from induced pluripotent stem cells (iPSC-RGCs) treated with H_2_0_2_. Increased expression of *SOD1* at 650uM H_2_0_2_ in iPSC-RGCs (Table 3a) and at 100uM H_2_0_2_ in hTM (Table 3b) indicates suitable concentrations of H_2_O_2_ for inducing oxidative stress conditions in these cell lines. Among all the genes tested, we found increased expression of *ADAM12, COL15A1, DNM3,* and *TRIM66* in both iPSC-RGCs and hTM cells treated with H_2_0_2_ when compared to untreated cells. Expression of *IL6, ANXA2, CXCR7,* and *PAK1* was downregulated in iPSC-RGCs, but upregulated in hTM after oxidative stress. *LMX1B* and *COL19A1* were overexpressed in iPSC-RGCs, while in hTM cells, *LMX1B* expression was suppressed after oxidative stress and *COL19A1* was undetectable. This differential expression of genes identified by our GWAS in iPSC-RGCs and hTM cells indicates their role in the maintenance of cellular homeostasis in RGCs and hTM cells under stress conditions.

**Table 3.** Evaluation of the expression of genes associated with POAG under oxidative stress condition, in (a) induced pluripotent stem cell derived retinal ganglion cells, and (b) and trabecular meshwork (TM) cells.

| Gene | Sample | Target gene average Ct ± SD | $\beta$ actin average Ct ± SD | ΔCt (Target gene average Ct - $\beta$ actin average Ct) ± SD | $\Delta\Delta$Ct (ΔCt treated- ΔCt untreated) ± SD | Fold difference in treated relative to untreated (CI) |
| --- | --- | --- | --- | --- | --- | --- |
| SOD1 | No H<sub>2</sub>O<sub>2</sub> (untreated) | 27.21±0.114 | 17.92±0.02 | 9.29±0.12 | 0±0.08 | 1(0.92-1.08) |
| | 325 μM H<sub>2</sub>O<sub>2</sub> (treated) | 27.41±0.05 | 16.99±0.01 | 10.42±0.05 | 1.13±0.03 | 0.46(0.44-0.47) |
| | 650 μM H<sub>2</sub>O<sub>2</sub> (treated) | 30.57±0.06 | 24.19±0.15 | 6.38±0.16 | -2.91±0.11 | **7.49**(6.71-8.36) |
| NRF2 | No H<sub>2</sub>O<sub>2</sub> (untreated) | 23.48±0.11 | 17.92±0.02 | 5.56±0.11 | 0±0.08 | 1(0.92-1.08) |
| | 325 μM H<sub>2</sub>O<sub>2</sub> (treated) | 23.56±0.13 | 16.99±0.01 | 6.57±0.13 | 1.01±0.09 | 0.5(0.45-0.54) |
| | 650 μM H<sub>2</sub>O<sub>2</sub> (treated) | 30.28±0.333 | 24.19±0.15 | 6.09±0.36 | 0.53±0.26 | 0.69(0.54-0.89) |
| IL6 | No H<sub>2</sub>O<sub>2</sub> (untreated) | 29.09±0.065 | 17.9±0.15 | 11.18±0.17 | 0±0.12 | 1(0.89-1.12) |
| | 325 μM H<sub>2</sub>O<sub>2</sub> (treated) | 27.76±0.02 | 16.32±0 | 11.44±0.02 | 0.25±0.01 | 0.84(0.48-0.82) |
| | 650 μM H<sub>2</sub>O<sub>2</sub> (treated) | 37.55±0.64 | 26.32±0.04 | 11.23±0.64 | 0.04±0.45 | 0.97(0.93-1.49) |
| ANAX2 | No H<sub>2</sub>O<sub>2</sub> (untreated) | 18.96±0.057 | 17.68±0.36 | 1.28±0.37 | 0±0.26 | 1(0.78-1.29) |
| | 325 μM H<sub>2</sub>O<sub>2</sub> (treated) | 18.71±0.09 | 16.04±0.16 | 2.67±0.18 | 1.39±0.13 | 0.38(0.48-0.82) |
| | 650 μM H<sub>2</sub>O<sub>2</sub> (treated) | 24.63±0.01 | 22.24±0.09 | 2.39±0.09 | 1.11±0.07 | 0.46(0.93-1.49) |
| ADAM12 | No H<sub>2</sub>O<sub>2</sub> (untreated) | 25.67±0.1 | 17.68±0.36 | 7.99±0.37 | 0±0.26 | 1(0.77-1.3) |
| | 325 μM H<sub>2</sub>O<sub>2</sub> (treated) | 22.39±0.05 | 16.04±0.16 | 6.35±0.17 | -1.64±0.12 | **3.12**(2.77-3.5) |
| | 650 μM H<sub>2</sub>O<sub>2</sub> (treated) | 30.76±0.12 | 22.24±0.09 | 8.52±0.15 | 0.52±0.11 | 0.7(0.63-0.77) |
| COL15A1 | No H<sub>2</sub>O<sub>2</sub> (untreated) | 31.37±0.01 | 17.68±0.36 | 13.69±0.36 | 0±0.26 | 1(0.78-1.28) |
| | 325 μM H<sub>2</sub>O<sub>2</sub> (treated) | 31.02±0.43 | 16.04±0.16 | 14.98±0.46 | 1.29±0.33 | 0.41(0.3-0.56) |
| | 650 μM H<sub>2</sub>O<sub>2</sub> (treated) | 31.8±0.13 | 22.24±0.09 | 9.55±0.16 | -4.14±0.11 | **17.6**(15.76-19.66) |
| COL19A1 | No H<sub>2</sub>O<sub>2</sub> (untreated) | 34.59±0.67 | 17.68±0.36 | 16.91±0.76 | 0±0.54 | 1(0.59-1.7) |
| | 325 μM H<sub>2</sub>O<sub>2</sub> (treated) | 32.12±0.012 | 16.04±0.16 | 16.08±0.16 | -0.83±0.11 | 1.77(1.59-1.98) |
| | 650 μM H<sub>2</sub>O<sub>2</sub> (treated) | 35.81±0 | 22.24±0.09 | 13.57±0.09 | -3.34±0.07 | **10.13**(9.49-10.8) |
| CXCR7 | No H<sub>2</sub>O<sub>2</sub> (untreated) | 20.93±0.032 | 17.68±0.36 | 3.25±0.36 | 0±0.26 | 1(0.78-1.29) |
| | 325 μM H<sub>2</sub>O<sub>2</sub> (treated) | 20.87±0.09 | 16.04±0.16 | 4.83±0.19 | 1.58±0.13 | 0.33(0.29-0.38) |
| | 650 μM H<sub>2</sub>O<sub>2</sub> (treated) | 26.42±0.06 | 22.24±0.09 | 4.17±0.11 | 0.92±0.08 | 0.53(0.49-0.57) |
| *DNM3* | No H<sub>2</sub>O<sub>2</sub><br>(untreated) | 25.56±0.237 | 17.68±0.36 | 7.88±0.43 | 0±0.31 | 1(0.74-1.35) |
|  | 325 µM H<sub>2</sub>O<sub>2</sub><br>(treated) | 26.3±0.37 | 16.04±0.16 | 10.26±0.4 | 2.38±0.28 | 0.19(0.15-0.25) |
|  | 650 µM H<sub>2</sub>O<sub>2</sub><br>(treated) | 27.46±0.05 | 22.24±0.09 | 5.22±0.1 | -2.66±0.07 | **6.33**(5.89-6.8) |
| *LMX1B* | No H<sub>2</sub>O<sub>2</sub><br>(untreated) | 32.32±0.181 | 17.68±0.36 | 14.64±0.4 | 0±0.29 | 1(0.76-1.32) |
|  | 325 µM H<sub>2</sub>O<sub>2</sub><br>(treated) | 32.18±0.22 | 16.04±0.16 | 16.14±0.27 | 1.5±0.19 | 0.35(0.29-0.43) |
|  | 650 µM H<sub>2</sub>O<sub>2</sub><br>(treated) | 35.84±0.25 | 22.24±0.09 | 13.6±0.27 | -1.04±0.19 | **2.06**(1.71-2.49) |
| *PAK1* | No H<sub>2</sub>O<sub>2</sub><br>(untreated) | 24.06±0.155 | 17.68±0.36 | 6.38±0.39 | 0±0.28 | 1(0.76-1.31) |
|  | 325 µM H<sub>2</sub>O<sub>2</sub><br>(treated) | 24.2±0.02 | 16.04±0.16 | 8.16±0.16 | 1.78±0.11 | 0.29(0.26-0.33) |
|  | 650 µM H<sub>2</sub>O<sub>2</sub><br>(treated) | 29.06±0.02 | 22.24±0.09 | 6.82±0.1 | 0.44±0.07 | 0.74(0.69-0.79) |
| *TRIM66* | No H<sub>2</sub>O<sub>2</sub><br>(untreated) | 27.95±0.02 | 17.68±0.36 | 10.27±0.36 | 0±0.26 | 1(0.78-1.28) |
|  | 325 µM H<sub>2</sub>O<sub>2</sub><br>(treated) | 27.28±0.04 | 16.04±0.16 | 11.24±0.16 | 0.98±0.12 | 0.51(0.45-0.57) |
|  | 650 µM H<sub>2</sub>O<sub>2</sub><br>(treated) | 30.91±0.083 | 22.24±0.09 | 8.66±0.12 | -1.6±0.09 | **3.04**(2.79-3.31) |

| Genes | Sample | Target gene<br>average Ct ±<br>SD | β actin<br>average<br>Ct ± SD | ΔCt (Target<br>gene average<br>Ct - β actin<br>average Ct) ±<br>SD | ΔΔCt (ΔCt<br>treated- ΔCt<br>untreated) ±<br>SD | Fold difference<br>in treated<br>relative to<br>untreated (CI) |
| --- | --- | --- | --- | --- | --- | --- |
| *SOD1* | No H<sub>2</sub>O<sub>2</sub><br>(untreated) | 24.33±0.063 | 16.13±0.14 | 8.2±0.15 | 0±0.11 | 1(0.9-1.11) |
|  | 100 µM H<sub>2</sub>O<sub>2</sub><br>(treated) | 25.34±0.05 | 18.29±0.09 | 7.05±0.1 | -1.15±0.07 | 2.21(2.06-2.37) |
| *NRF2* | No H<sub>2</sub>O<sub>2</sub><br>(untreated) | 20.44±0.376 | 16.13±0.14 | 4.31±0.4 | 0±0.28 | 1(0.76-1.32) |
|  | 100 µM H<sub>2</sub>O<sub>2</sub><br>(treated) | 22.58±0.11 | 18.29±0.09 | 4.3±0.14 | -0.01±0.1 | 1.01(0.91-1.11) |
| *IL6* | No H<sub>2</sub>O<sub>2</sub><br>(untreated) | 26.22±0.061 | 16.11±0.01 | 10.11±0.06 | 0±0.04 | 1(0.96-1.04) |
|  | 100 µM H<sub>2</sub>O<sub>2</sub><br>(treated) | 29.5±0.27 | 20.83±0.07 | 8.67±0.28 | -1.44±0.2 | **2.71**(2.23-3.3) |
| *ANXA2* | No H<sub>2</sub>O<sub>2</sub><br>(untreated) | 19.47±0.034 | 16.82±0.17 | 2.68±0.17 | 0±0.17 | 1(0.89-1.12) |
|  | 100 µM H<sub>2</sub>O<sub>2</sub><br>(treated) | 22.76±0.072 | 22.64±0.01 | 0.17±0.07 | -2.5±0.07 | **5.67**(5.39-5.97) |
| *ADAM12* | No H<sub>2</sub>O<sub>2</sub><br>(untreated) | 24.13±0.11 | 16.82±0.17 | 7.31±0.2 | 0±0.14 | 1(0.87-1.15) |
|  | 100 µM H<sub>2</sub>O<sub>2</sub><br>(treated) | 29.47±0.16 | 22.64±0.01 | 6.83±0.16 | -0.48±0.11 | 1.4(1.25-1.56) |
| *COL15A1* | No H2O2<br>(untreated) | 32.55±0.003 | 15.73±0.17 | 15.73±0.17 | 0±0.12 | 1(0.89-1.12) |
|  | 100 µM H2O2<br>(treated) | 35.87±0.01 | 13.23±0.02 | 13.23±0.02 | -2.5±0.01 | **5.64**(5.56-5.72) |
| *CXCR7* | No H2O2<br>(untreated) | 26.68±0.227 | 16.82±0.17 | 9.86±0.28 | 0±0.2 | 1(0.82-1.21) |
|  | 100 µM H2O2<br>(treated) | 27.62±0.12 | 22.64±0.01 | 4.98±0.12 | -4.88±0.08 | **29.35**(27.06-31.84) |
| *DNM3* | No H2O2<br>(untreated) | 31.75±0.77 | 16.82±0.17 | 14.93±0.79 | 0±0.56 | 1(0.58-1.73) |
|  | 100 µM H2O2<br>(treated) | 33.8±0.28 | 22.64±0.01 | 11.16±0.28 | -3.76±0.2 | **13.58**(11.16-16.53) |
| *LMX1B* | No H2O2<br>(untreated) | 23.7±0.176 | 16.82±0.17 | 6.88±0.24 | 0±0.17 | 1(0.85-1.18) |
|  | 100 µM H2O2<br>(treated) | 30.39±0.02 | 22.64±0.01 | 7.75±0.03 | 0.87±0.02 | 0.55(0.54-0.56) |
| *PAK1* | No H2O2<br>(untreated) | 25.87±0.094 | 16.82±0.17 | 9.05±0.19 | 0±0.13 | 1(0.88-1.14) |
|  | 100 µM H2O2<br>(treated) | 27.66±0.07 | 22.64±0.01 | 5.02±0.07 | -4.02±0.05 | **16.24**(15.44-17.09) |
| *TRIM66* | No H2O2<br>(untreated) | 27.66±0.08 | 16.82±0.17 | 10.84±0.18 | 0±0.13 | 1(0.88-1.14) |
|  | 100 µM H2O2<br>(treated) | 31.95±0 | 22.64±0.01 | 9.31±0.01 | -1.52±0.01 | **2.88**(2.85-2.91) |

Fold change $\geq 2$ considered significantly differentially expressed and highlighted in bold. Standard deviation represented as SD

### Pathway analyses highlights heterogeneous nature of POAG

Pathway analysis using ingenuity pathway analysis (IPA) (p-value<0.05, Table 4) showed that matrix metalloproteinase inhibition and tight junction signaling pathways were significant for CDR associated genes, while remodeling of adherens junction was moderately significant for IOP associated genes. Receptor signaling pathways such as DNA damage-induced signaling, Amyotrophic lateral sclerosis (ALS) signaling, synaptogenesis, Ephrin signaling, and genes associated with Pyridoxal 5’ pathways that prevent ganglion cell loss due to ischemic retinal injury were significant for genes associated with MD and PSD in our study.^62^ Pathways identified by IPA analyses were not shared among any quantitative traits. Pathway analysis for genes obtained from MAGMA analysis identified genes in tight junction signaling, retinol biosynthesis, and coagulation as significant for the development of POAG (Table 4).

**Table 4:** Pathways analysis of POAG-associated genes for various endophenotypes and of genes from MAGMA analysis using ingenuity pathways analysis.

| Phenotype | Ingenuity Canonical Pathways | p-value | genes |
| --- | --- | --- | --- |
| CDR | Inhibition of Matrix Metalloproteases* | 0.0069 | *ADAM12* |
|  | Tight Junction Signaling* | 0.035 | *MPDZ* |
|  | Axonal Guidance Signaling | 0.092 | *ADAM12* |
| IOP & IOP max | Remodeling of Epithelial Adherens Junctions | 0.05 | *DNM3* |
|  | FAK Signaling | 0.071 | *TNS1* |
| MD | Pyridoxal 5'-phosphate Salvage Pathway* | 0.006 | *CDK8,PAK1* |
|  | Agrin Interactions at Neuromuscular Junction* | 0.008 | *ERBB4,PAK1* |
| | DNA damage-induced 14-3-3σ Signaling* | 0.025 | *ATR* |
| PSD | Amyotrophic Lateral Sclerosis Signaling* | 0.01 | *GRIN2C,PAK1* |
|  | Ephrin Receptor Signaling* | 0.0034 | *GRIN2C,PAK1* |
|  | Synaptogenesis Signaling Pathway* | 0.01 | *GRIN2C,PAK1* |
|  | Retinol Biosynthesis* | 0.034 | *CES4A* |
|  | Role of Oct4 in Mammalian Embryonic Stem Cell Pluripotency | 0.055 | *PHB* |
|  | Synaptogenesis Signaling Pathway | 0.061 | *GOSR1,STXBP3* |
|  | Semaphorin Signaling in Neurons | 0.077 | *CRMP1* |

P-value >0.05 is significant. CDR=cup-to-disc ratio, IOP=intraocular pressure in mmHg, MD=mean deviation, PSD=pattern standard deviation

## Discussion

The majority of the genetic studies on POAG to date have been conducted in individuals of European and Asian ancestry. Our study recruited, genotyped, and deeply phenotyped the largest cohort of African Americans with POAG recruited from a single city, providing novel information on disease biology for this most affected population. The GWAS study in 5950 African Americans implicated four variants with higher risk of POAG, as well as 260 variants in 127 genomic regions with glaucoma-related endophenotypes.

In the case-control GWAS analysis, we identified one variant (rs112369934) located near the *TRIM66* gene that reached genome-wide significance. *TRIM66* belongs to the Tripartite motif *(Trim)* family of genes that are highly represented in the retinal progenitor cells (RPCs) in the developing mouse retina.^63^ We also used MAGMA software analysis to determine gene and gene-set based associations, which employs a multiple regression model.^64^ MAGMA analysis suggested *CLCF1* as a significant gene for POAG in sex-combined case-control analysis. *CLCF1* encodes an inflammatory cytokine, cardiotrophin like cytokine factor 1 protein (CLCF1). CLCF1 is a heterodimeric neurotropic cytokine that may play a role in the maintenance of RGC health in POAG. CLCF1 demonstrates both growth factor and neuroprotective effects on RGCs,^65^ and its expression has been associated with response to stress as observed in glaucoma.^66^

The GWAS analysis also confirmed the association of several previously associated POAG variants in other populations, including *CDKN2B-AS1, ARHGEF12,* and *TMCO1,* supporting the likelihood of their contribution to POAG. However, many previously associated loci identified in other ancestral populations did not replicate in our study. This is likely due to genetic heterogeneity across different ancestral groups. Because our study is the largest African American dataset available to date, no comparable ancestry-matched dataset was available to replicate these novel findings.

The genome-wide quantitative trait association studies of seven distinct POAG phenotypic features identified loci in 127 genomic regions. Further functional studies, tissue localization, and loss-of-function studies are necessary to fully understand the role of these genes in POAG. The *CXCR7* gene associated with CDR is a chemokine receptor that controls the chemokine network. In glioma cells, *CXCR7* transduces signals via MEK/ERK pathway, mediating resistance to apoptosis by promoting cell growth and survival.^67^ The *MPDZ* gene, also associated with CDR, encodes a multiple PDZ domain crumbs cell polarity complex protein. Novel variants in this gene were previously reported in a multi-ethnic Asian GWAS study to be associated with CCT but not with CDR.^68^ Increased expression of *ADAM12-L* was reported in TM cells in response to oxidative stress resulting from elevated IOP, causing RGC degeneration and affecting the CDR.^69^

IOP baseline and IOP maximum traits were both associated with the *COL15A1* gene, which encodes the alpha chain of type XV collagen. A commonly reported variant in *COL15A1,* R163H is reported to influence the age of onset of POAG.^70^ RNA *in situ* hybridization of mouse eyes showed that *Col15a1* was expressed in multiple ocular structures including the ciliary body, astrocytes of the ON, and cells in the RGC layer.^70^

For the genes associated with visual fields, previous studies have shown that *Lmx1b* is needed for the formation of the TM and maintenance of corneal transparency in the mouse anterior segment.^71^ A dominant negative mutation of mouse *Lmx1b (Lmx1b^Icst/+^*) also causes glaucoma and is semi-lethal.^72^ The *COL19A1* gene encodes the alpha chain of type XIX collagen, a member of the FACIT collagen family (fibril-associated collagens with interrupted helices). Decreased expression of *Col19A1* (1.9 fold) was observed in a microarray of an 8-month old DBA/2J mouse retina when compared to 3-month-old retina. This suggests the encoded protein may play a role in the maintenance of the cytoskeletal matrix as a result of prolonged IOP elevation in the older mice, which can result in ganglion cell loss.^73^ The *PAK1* gene encodes a family member of serine/threonine p21-activating kinases, known as PAK proteins.^74^ PAK1 is inactivated by oxidative stress conditions and oxidative stress-induced down-regulation of PAK1 activity could be involved in the loss of mesencephalic dopaminergic (DA) neurons through modulation of neuronal death in Parkinson’s disease,^75^ suggesting that the PAK1 protein may have a role in neuroprotection.

Our study identified genes that have significantly altered expression in TM cells *(ADAM12, COL19A1*, *DNM3,* and *TRIM66*), suggesting that these genes may play a role in pathways leading to TM cell damage or altered homeostasis. Inflammatory pathway genes identified in our study including *IL6*, *ANXA2*, *COL15A1*, *CXCR7*, *DNM3,* and *PAK1* exhibit gene transcripts that were significantly overexpressed in iPSC-RGCs treated with hydrogen peroxide, indicating that these genes may be involved in maintaining RGC health and could play a role in glaucomatous neurodegeneration. These findings are also supported by our IPA analysis performed using the implicated genes from this study.

As this is the first large effort to study the genetic basis of POAG in a deeply phenotyped African American cohort, the findings we present here must be replicated in future cohorts. Notably, associations from the glaucoma binary (case-control) phenotype are different than the associations with quantitative traits in cases and controls, likely reflecting heterogeneity in the underlying genetic etiologies of glaucoma. A recently published GWAS on individuals of African ancestry identified a variant on chromosome 4 (rs59892895) linked with POAG case-control status.^41^ This association did not replicate in our study (p-value=0.335). Additionally, our genome-wide significant SNP from case-control analyses (rs112369934) was not available in summary statistics in this published study. Differences between the GWAS findings in other POAG cohorts should be examined carefully, and follow-up analyses should consider genetic ancestry when interpreting these differences. Our study investigated African Americans, which is a more admixed population compared to the African descent individuals.^76,77^ Additional analyses will help validate the biological implications of the variants discovered from the POAAGG study.

Enviornmental factors can play a confounding role in power to detect associations. Strengths of this study include the recruitment of African Americans solely from Philadelphia, allowing us to control for differences in environment and socioeconomic factors, possibly increasing statistical power to detect GWAS associations. This study also employed the MEGA array, which had custom content designed through selective whole-genome sequencing, maximizing coverage of variants relevant for genomic analysis of admixed populations. We carefully ascertained endophenotypes from each subject, which is crucial to prevent residual confounding effects of unmeasured phenotypes within association studies.^78^

A limitation of this study is that the genetic variation in African Americans is greater than that observed in European Americans,^79^ limiting power to detect SNP effects in a GWAS analysis and necessitating a larger cohort to detect significant associations. However, the POAAGG study as a whole presents a unique opportunity to examine an admixed population, potentially leading to novel discoveries that may not be apparent in studies that focus on samples with homogeneous ancestry. Another limitation is the lack of replication of our significant associations from the quantitative traits; however, to our knowledge, no African ancestry dataset with similar endophenotypes is available yet.

In conclusion, this study greatly expands knowledge of the genetic underpinnings of POAG in African Americans. Future analyses will help validate the biological implications of the variants discovered from this study, and have potential application to screening and treating this overaffected population.

## Methods

### Subjects

#### Eligibility Criteria

The POAAGG study population consists of self-identified blacks (African American, African descent, or African Caribbean), aged 35 years or older. Exclusion criteria included a history of narrow angle, closed angle, neovascular, mixed mechanism, or pseudoexfoliation glaucoma; history of glaucoma secondary to eye surgery or secondary to severe ocular trauma; history of iritis, uveitis, or iridocyclitis; presence of Grave’s disease with ocular manifestations, vascular occlusion causing neovascularization of the iris, optic nerve atrophy from other diagnoses, or advanced proliferative diabetic retinopathy. All subjects provided informed written consent, in accordance with the tenets of the Declaration of Helsinki, under IRB-approved protocols.

#### Study Procedures

Study participants were identified from within all comprehensive and subspecialty ophthalmology clinics at the University of Pennsylvania (Scheie Eye Institute, Perelman Center for Advanced Medicine, Mercy Fitzgerald Hospital, VA Hospital); Lewis Katz School of Medicine at Temple University; Drexel University College of Medicine; and a private practice (Windell Murphy, MD). The clinical examination included an onsite ophthalmic exam and interview. Examination data were entered directly into the Research Electronic Data Capture (REDCap) database utilized at University of Pennsylvania (UPenn).^80^ Data collection was standardized and included a series of measurements taken analyzed glaucoma specialists.

The full onsite exam included: (1) verification of name, age, date of birth, street address, gender, and informed consent with signature; (2) completion of a health and family history questionnaire in-clinic including evaluation of diabetes and/or high blood pressure diagnoses, ocular surgical history, family history of glaucoma, and tobacco and alcohol use (confirmed in medical record when possible); (3) evaluation of height and weight; (4) explanation of procedure for blood or saliva collection for DNA analysis; (5) visual acuity measured using Snellen chart at 20 feet; (6) automated refraction with a Reichert Phoropter RS Automatic Refractor (Reichert Technologies, Depew, NY) if the presented visual a acuity was not 20/20 in either eye, followed by manual refraction; (7) near vision assessed using the Snellen chart at near with the participant’s current reading prescription; (8) intraocular pressure measured with a Goldmann applanation tonometer; (9) anterior and posterior segment examinations by slit lamp with a 90-diopter lens; (10) gonioscopy; (11) central corneal thickness and axial length measurements assessed with an ultrasonic A-scan/pachymeter DGH 4000B SBH IOL Computation module (DGH Tech Inc., Exton, PA); (12) visual field test utilizing the Humphrey Automated Field Analyzer (Standard 24-2 Swedish interactive thresholding algorithm); (13) stereo disc photos and fundus photography utilizing the Topcon TRC 50EX Retinal Camera (Topcon Corp. of America, Paramus, NJ); and (14) optical coherence tomography (OCT) using either Cirrus or Stratus OCT (Carl Zeiss Meditec, Dublin, CA) or iVue (Optovue, Fremont, CA). The outcomes of the procedures and all diagnoses were discussed with the patient at the conclusion of the examination.

#### Phenotyping

Cases and controls were selected from the UPenn patient population based on the following criteria: Primary open-angle glaucoma (POAG) cases were defined as having an open iridocorneal angle and: (1) characteristic glaucomatous optic nerve findings in one or both eyes consisting of at least one of the following: notching, neuroretinal rim thinning, excavation, or a nerve fiber layer defect; (2) characteristic visual field defects on two consecutive reliable visual field tests in at least one eye, which were consistent with the observed optic nerve defects in that eye, as determined by fellowship-trained glaucoma specialists; and (3) all secondary causes of glaucoma excluded. Normal controls were defined as subjects older than 35, without: (1) high myopia (greater than – 8.00 diopters); (2) high presbyopia (+8.00 diopters); (3) abnormal visual field; (4) intraocular pressure greater than 21 mmHg; (5) neuroretinal rim thinning, excavation, notching or nerve fiber layer defects; (6) optic nerveasymmetry; or (7) a cup-to-disc ratio difference between eyes greater than 0.2. A preliminary masked concordance study among glaucoma specialists at three institutions confirmed a 97% concordance rate in diagnosis of 120 glaucoma cases and controls.^54^ We collected the below phenotypic measures.

1. Central corneal thickness (300–800 microns): measure of the thickness of the cornea
2. Cup-to-disc ratio (>0–1, decimal fraction): measure of extent of cupping of the optic nerve in relation to the disc, with a higher ratio indicating greater degeneration of the nerve
3. Intraocular pressure (>0–70 mmHg): the fluid pressure within the eye, which often (but not always) is elevated in eyes with POAG. Maximum IOP was defined as the highest documented IOP value for each patient.
4. Mean deviation (−35-+20 dB): quantitative output from a Humphrey visual field test that represents the average of a patient’s deviation from the visual sensitivity of age-matched controls at all test locations; lower values indicate greater differences from controls and worsening visual fields
5. Pattern standard deviation (0–20 dB): quantitative output from a Humphrey visual field test that quantifies irregularities in the field, with larger values signifying non-uniform sensitivity loss
6. Retinal nerve fiber layer (30–130 microns): mean thickness of the nerve fiber layer, which contains fibers expanding into the optic nerve; lower values typically indicate loss of cells and increased severity of POAG
7. Visual acuity (−0.3–6.0, logMAR units): measure of the clarity of central vision using a Snellen or logMAR chart. All Snellen values were converted to logMAR units using a published formula.^81^ We used corrected VA for this analysis.

The distribution of available quantitative phenotypes used for the association analysis is shown in box plots in Extended Figure 2.

### Genotyping and Quality Control

#### Specimen Collection

Blood was collected by venipuncture in 10 milliliter purple top tubes with EDTA anticoagulant. These samples were frozen at –20 degrees prior to DNA isolation. For saliva collection, subjects were asked to refrain from smoking, drinking, or eating prior to donating specimens. Two milliliters of saliva per subject were collected in Oragene DISCOVER (OGR-500) self-collection kits (DNA Genotek, Canada). The saliva specimens were mixed with stabilizing reagent within the collection tubes per manufacturer’s instructions.

#### DNA Extraction and Quantitation

Genomic DNA for all enrolled subjects was extracted from peripheral blood or saliva. DNA was isolated from thawed blood samples using Gentra PureGene kits (Qiagen, Valencia, CA) and included the optional RNase treatment step. DNA from saliva samples was extracted using the prepIT.L2P reagent (cat # PT-L2P-5, DNA Genotek, Canada) and precipitated with ethanol according to manufacturer’s instructions. The saliva DNA samples were RNase treated by double digestion with RNase A and RNase T and re-precipitated using ethanol according to the manufacturer’s instructions. The concentrations of DNA from blood and saliva samples were determined using the fluorescence-based Quant iT dsDNA Board-Range assay kit (cat # Q-33130, Life Technologies, CA). Fluorescence was measured with a Tecan Infinite M 200 Pro multimode microplate reader (Tecan, NC).

#### Genotyping

A 25 μl aliquot of all samples with high-quality DNAs and case/control status were plated for array-based high throughput genotyping. These samples consisted of DNA extracted from blood (67.3%) and DNA isolated from saliva (32.7%), as described above. Prior to performing these studies, the UPenn and Illumina labs demonstrated a 99.996% concordance rate on this array between blood and saliva.^82^

Genotyping was attempted on 6525 DNA samples using the Multi-Ethnic Genotyping Array (MEGA)V2 (EX) consortium chip on the Infinium iSelect platform by Illumina FastTrack Services (Illumina, San Diego, CA). Standard MEGA array content was supplemented with 5000 SNPs from prior GWAS studies on POAG, common polymorphisms from POAG-associated genes, and variants detected from whole-genome sequencing of POAAGG subjects. At least one sample per 96-well microtiter plate was genotyped in duplicate for purposes of quality control. A total of 6406 samples passed genotyping. The genotype calls were generated using the Genome Studio genotyping module (GT). Cluster optimization and reproducibility analysis for paired samples were performed as per standard practices at Illumina FastTrack services.

#### Quality Control

Standard quality assurance/quality control (QA/QC) methods (Extended Table 1, 2) for sample and variant quality filtering were applied.^83^ Pre-imputation QC, including a comparison between reported gender and genetically informed sex and 95% sample and marker call rate filters, was run on 2,040,811 SNPs and 6,525samples. A total of 1,845, 896 variants and 6403 people passed preimputation quality control measures while 775,309 variants were ultimately used for imputations after phasing and strand checks.

#### Imputation

Imputation was carried out across the autosomal chromosomes using the full 1000 Genomes Project Phase 3 reference panel (N=2504).^84^ The study samples were imputed together using genotyped SNPs passing quality filters as described above, and were first phased using Eagle2 and then imputed using the Michigan Imputation Server v1.0.4 (Minimac 3).^85^ Association analyses were performed on all variants with an info score > 0.3.

### Statistical Analyses

#### Single Variant Association Testing

Case-control analyses were conducted on 5950 DNA samples (2589 cases and 3361 controls) to detect associations with POAG risk. Variants that failed QC based on missingness filter criteria as described in Extended Table 2 were excluded. No samples faild 99% misisngness call rate filter after imputation. Additionally, we removed 453 related individuals using a priority-based network approach on samplew with pi-hat >=0.25. Samples were removed in the following order of priority: samples related to multiple other samples in the dataset, controls, and samples with a high degree of missingness in quantitative traits.

Association analyses were adjusted for the first five principal components (PCs), age, and sex to prevent spurious association and inflation of test statistics due to population structure or confounding factors. The PCs were calculated by running Eigensoft SmartPCA on uncorrelated, QCed SNPs and projected onto an external set of 1000 Genomes samples^84^. The study samples fall along the gradient from the cluster of European to the cluster of African ancestry reference samples, consistent with the expected genetic ancestry from self-identified African American cohort (Extended Data Figure 1).

Single variant, binary association tests were performed genome-wide using a logistic model framework as implemented in the PLATO software package.^86^ Covariates of sex, age at study enrollment, and the first five PCs were included in the model as fixed effects for the entire POAG case control and VA logMAR analysis. Age at study enrollment and the first five PCs were included as covariates in the model for the sex-stratified GWAS.

Single variant, quantitative trait association tests were performed genome-wide using generalized estimating equations (GEE) as implemented in the gee R software package (https://CRAN.R-project.org/package=gee). This method adjusts for the inter-eye correlation for measures taken from both eyes of a subjects and subjects with either one or two eye measurements can be included in GEE analysis.^87,88^ As in the case-control analysis, covariates of sex, age at study enrollment and the first five PCs were included as fixed effects. Outcomes of baseline IOP (n=5282 samples), maximum IOP (n=5282 samples), baseline CDR (n=5049 samples), ‘latest’ central CCT (n=2616 samples), baseline visual acuity (n=5483 samples), baseline mean deviation (MD baseline, n=2213 samples), baseline pattern standard deviation (PSD baseline, n=2211 samples) and baseline retinal nerve fiber layer (RNFL, n=2239 samples) were tested in all individuals for whom measurements were available.

Samples passing QC with non-missing information for all covariates were included in the association tests. Calibration of the association tests was examined using QQ plots and the genomic control inflation factor,^89^ calculated as the ratio of the median observed test statistic to the median expected test statistic; a value of 1 indicates the observed and expected median test statistics are equivalent. Variants were considered statistically significant if the estimated score test p-value < 5×10^-8^. Additionally, odds ratios and standard errors were estimated for the four significant or suggestive variants from the case-control GWAS using the Wald test. Post-GWAS, we evaluated all results were evaluated using FUMA^90^ web-based tool. Gene based analyses using MAGMA was performed using FUMA, SNPs in our results were mapped to 18,991 genes. All genes below Bonferroni corrected p-value (0.05/18991 = 2.63e-06) were reported as significant.

#### In silico analyses for gene function in ocular tissue databases

To understand the functional relevance of genes associated with POAG in our study, we evaluated expression of the genes that contained associated credible set variants in human ocular tissues using a publicly available database: the OTDB (https://genome.uiowa.edu/otdb/).^58^ The OTDB consists of gene expression data for eye tissues from 20 normal human donors generated using Affymetrix Human Exon 1.0 ST arrays.^58^ The gene expression is described as Affymetrix Probe Logarithmic Intensity Error (PLIER) values where individual gene expression values are normalized to expression in other tissues.

We assessed the gene expression in mouse ocular tissues using a publicly available domain “Glaucoma Discovery Platform” (http://glaucomadb.jax.org/glaucoma).^59^ This platform provides an interactive way to analyze RNA sequencing data obtained from RGCs isolated from retina and optic nerve head of 9-month-old female D2 mouse (an age-dependent model of ocular hypertension/glaucoma), and *D2-Gpnmb^+^* mouse (that does not develop high IOP/glaucoma).^91^ In this study, four distinct groups of D2 and *D2-Gpnmb^+^* mice were compared based on the axonal degeneration, and gene expression patterns. The transcriptome of D2 group 1 is identical to the control strain (D2-*Gpnmb^+^*); and D2 groups 2–4 exhibit increasing levels of molecular changes relevant to axonal degeneration when compared to the control group.^92^

An interactive DATAGAN software queried the transcriptome data from D2 and *D2-Gpnmb^+^* mice to identify differentially expressed genes in these mouse tissues undergoing molecular changes.^59^ The output data from this software was represented as log_2_ fold change and *q*-value between different stages of D2 mice and D2. *Gpnmb+* (control) samples, with -*q*-Values < 0.05 considered to be significant. We compared expression profiles of genes associated variants reaching significance in our GWAS study among D2 groups and control group.^59^ Among the genes implicated in the 95% credible set of variants, a few did not have mouse homologs or available data (e.g., *LOC105378189, LOC145783, LMX1B,* and *ZNF280D*).

#### TM and iPSC-RGC cultures

The primary TM cells were commercially obtained from Sciencell, CA, USA (Cat#6590) and also as a gift from Dr Markus Kuehn, University of Iowa.^42^ We validated the authenticity of the TM cells by performing MYOC induction experiments with dexamethasone as described previously.^42^ We used Passage 2 (P2) to Passage 4 (P4) primary TM cultures for all our experiments. The RGCs for our studies were derived using small molecules to inhibit BMP, TGF-beta (SMAD) and Wnt signaling to differentiate iPSCs into retinal ganglion cells (RGCs). The generation and characterization of these cells employed a slight modification of protocol from Teotia etal., 2017^93^ and is described in Chavali *et al.,* **doi:**(article in revision).

#### Evaluating oxidative stress in TM and iPSC-RGCs

The primary TM cells and induced pluripotent stem cell derived retinal ganglion cells (RGCs) were incubated with increased amounts of H_2_O_2_ overnight before replacing the cultures with complete media. The cells were collected 24 hours after the H_2_O_2_ treatment and levels of few selected gene transcripts form the our GWAS using quantitative RT-PCR and gene expression primers (Supplementary Table) and following previously published protocols.^94–96^ The relative gene expression was compared against control (no treatment) to obtain normalized gene expression. Expression levels (±SEM) were calculated by analyzing at least three independent samples with replicate reactions and presented on an arbitrary scale that represents the expression of relevant genes compared to the housekeeping gene ACTB. Increase in *Superoxide Dismutase 1(SOD1)* expression levels served as a positive controls for oxidative stress in the same set of samples.

#### Ingenuity Pathway Analysis (IPA)

All the SNPs in genes identified in our case-control association study and endophenotype association study were uploaded into Qiagen’s IPA system for core analysis. They were then collapsed with the molecular network found in databases from the brain, neurological tissues, and tissues from eye in the Ingenuity Pathway Knowledge Base (IPKB).^97,98^ IPA was performed to identify canonical pathways, diseases and functions, and gene networks that are most significant to genome-wide association findings from our study, and to categorize differentially expressed genes in specific diseases and their possible functions. The IPA analysis summary includes the associated cellular and molecular functions and development pathways with the predicted p-values of significance. The relevance of associated genes from our POAAGG study are discussed.

## Data availability

Genotype data of POAAGG participants are available from the database of Genotypes and Phenotypes (dbGaP) under accession phs001312.

## Notes

<u>Financial Support</u>: This work was supported by the National Eye Institute, Bethesda, Maryland (grant #1RO1EY023557-01) and Vision Research Core Grant (P30 EY001583). Funds also come from the F.M. Kirby Foundation, Research to Prevent Blindness, The UPenn Hospital Board of Women Visitors, and The Paul and Evanina Bell Mackall Foundation Trust. The Ophthalmology Department at the Perelman School of Medicine and the VA Hospital in Philadelphia, PA also provided support. The sponsor or funding organization had no role in the design or conduct of this research.

