## Supplement Data for "Genome wide-association study identifies novel loci in the Primary Open-Angle African American Glaucoma Genetics (POAAGG) study"

### Extended Data

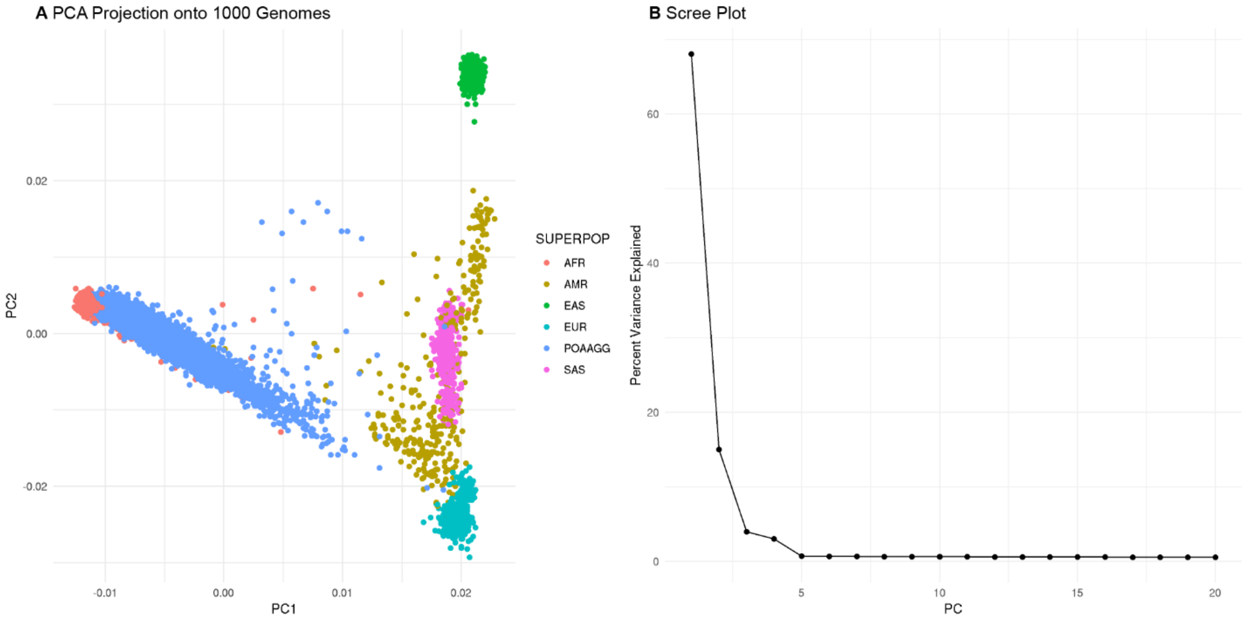


**Extended Data Fig. 1**: **PCA Results with 1000 Genomes Reference Samples. (a)** The first and second principal components from PCA analysis, including 5950 POAAGG study samples and 2535 1000 Genomes reference samples, colored by ancestry groups, and **(b)** the percentage variance explained by each eigenvector is displayed in the axis labels of the scree plot.

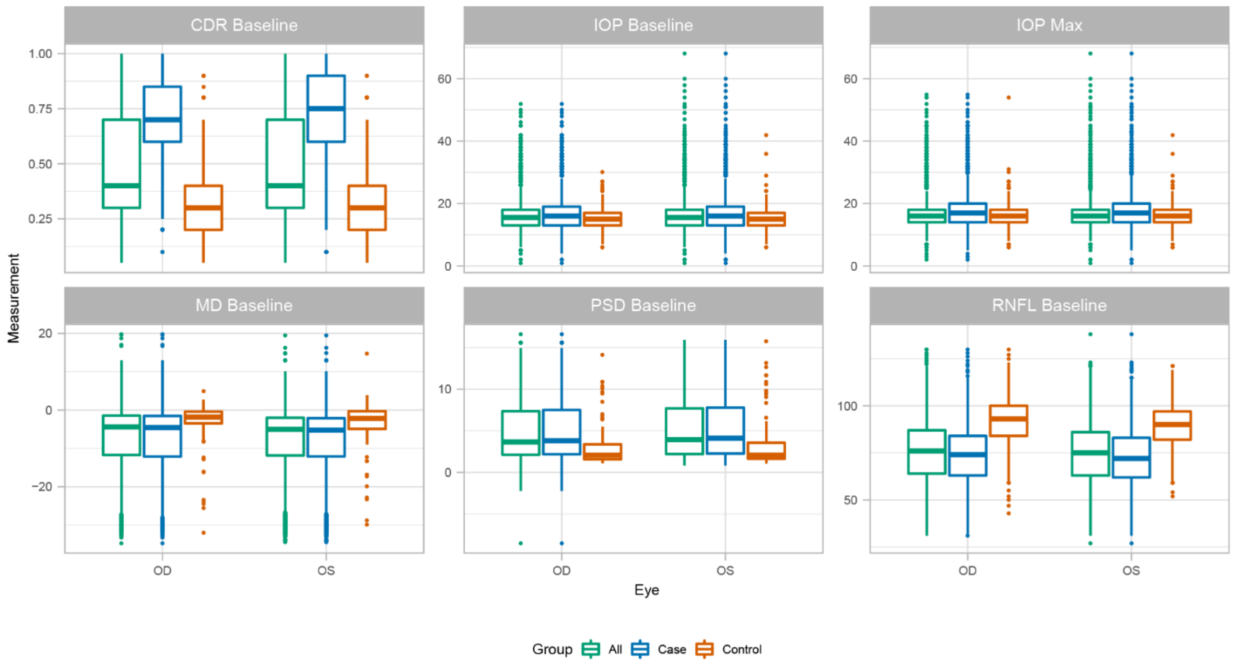


**Extended Data Fig. 2: Box plots representing summary of quantitative traits.** These plots include all subjects (yellow), cases only (blue), and controls only (green). The summary for both right (OD) and left (OS) eye measurements is represented in these plots.

**Extended Data Table 1**: Summary of sample quality control for genotyping.

| Sample QC step | N |
|---|---|
| Raw Data | 6525 |
| Sex Mismatch | 6406 |
| 95% Sample Call Rate | 6403 |
| Remove relateds | 5950 |

QC=quality control

**Extended Data Table 2:** Summary of variant quality control for genotyping.

| Stage | Filter | Number of SNPs |
|---|---|---|
| Pre-Imputation | Raw Genotype | 2,040,811 |
| | 95% Marker Call Rate | 1,845,896 |
| | Strand check | 803,835 |
| | Eagle2 Phasing | 775,309 |
| Post-Imputation | Raw Imputed | 47,118,013 |
| | Info Score > 0.3 | 33,073,928 |
| | 99% Marker Call Rate | 33,073,928 |
| | Minor Allele Frequency > 1% | 15,866,666 |

**Extended Data Table 3: All genome-wide significant variants from quantitative trait analyses.**

| Phenotype | Chr.Pos | Mapped/Nearest Gene | rsID | p.value | Beta | 95% CI | MAF |
|---|---|---|---|---|---|---|---|
| CDR | 10:127738557_G | ADAM12 | | 2.89E-08 | 0.03 | 0.02 - 0.03 | 0.411 |
| CDR | 11:3511587_C | RPS3AP39 | rs11027638 | 1.47E-08 | -0.05 | -0.06 - -0.03 | 0.088 |
| CDR | 11:3513525_T | RPS3AP39 | rs148849965 | 1.12E-08 | -0.05 | -0.07 - -0.03 | 0.063 |
| CDR | 11:3514393_G | RPS3AP39 | rs191137683 | 1.41E-10 | -0.06 | -0.07 - -0.04 | 0.074 |
| CDR | 11:3515761_G | RPS3AP39 | rs6578377 | 9.20E-11 | -0.06 | -0.07 - -0.04 | 0.074 |
| CDR | 11:3515991_C | RPS3AP39 | rs150613671 | 1.19E-10 | -0.06 | -0.07 - -0.04 | 0.074 |
| CDR | 11:3516254_C | | rs545312069 | 9.44E-11 | -0.06 | -0.07 - -0.04 | 0.074 |
| CDR | 11:3517193_T | RPS3AP39 | rs7934188 | 5.37E-11 | -0.06 | -0.07 - -0.04 | 0.073 |
| CDR | 11:3519141_T | RPS3AP39 | rs112885003 | 4.65E-10 | -0.05 | -0.07 - -0.04 | 0.069 |
| CDR | 11:3519378_T | RPS3AP39 | rs149215469 | 5.50E-11 | -0.06 | -0.07 - -0.04 | 0.074 |
| CDR | 11:3519418_A | RPS3AP39 | rs143254765 | 4.02E-09 | -0.05 | -0.06 - -0.03 | 0.088 |
| CDR | 11:3520545_T | RPS3AP39 | rs11027695 | 1.09E-09 | -0.05 | -0.06 - -0.03 | 0.093 |
| CDR | 11:3521989_T | RPS3AP39 | rs7110375 | 1.83E-13 | -0.06 | -0.08 - -0.05 | 0.068 |
| CDR | 11:3522217_C | RPS3AP39 | rs7126349 | 1.99E-11 | -0.06 | -0.07 - -0.04 | 0.072 |
| CDR | 11:3522501_T | RPS3AP39 | rs112051853 | 1.99E-11 | -0.06 | -0.07 - -0.04 | 0.072 |
| CDR | 11:3523069_G | RPS3AP39 | rs6578381 | 3.49E-11 | -0.06 | -0.07 - -0.04 | 0.072 |
| CDR | 11:3523100_T | RPS3AP39 | rs6578382 | 3.49E-11 | -0.06 | -0.07 - -0.04 | 0.072 |
| CDR | 11:3567134_A | RPS3AP39 | rs146887003 | 1.70E-08 | -0.08 | -0.1 - -0.05 | 0.027 |
| CDR | 11:3573251_T | RPS3AP39 | rs79280292 | 5.98E-10 | -0.07 | -0.1 - -0.05 | 0.035 |
| CDR | 11:3578116_C | RPS3AP39 | rs11028083 | 2.30E-09 | -0.07 | -0.09 - -0.05 | 0.036 |
| CDR | 13:52230897_G | WDFY2 | rs144971368 | 2.16E-08 | -0.1 | -0.13 - -0.06 | 0.011 |
| CDR | 1:48359572_G | TRABD2B | rs74774657 | 1.82E-08 | -0.09 | -0.12 - -0.06 | 0.013 |
| CDR | 15:22515589_T | MIR1268A | rs552375943 | 1.30E-10 | -0.07 | -0.1 - -0.05 | 0.039 |
| CDR | 19:55269926_C | KIR2DL3 | rs146547954 | 8.44E-15 | -0.07 | -0.09 - -0.05 | 0.066 |
| CDR | 19:55272110_T | KIR2DL1 | rs184422553 | 2.18E-15 | -0.07 | -0.09 - -0.06 | 0.063 |
| CDR | 2:237653539_G | CXCR7 | rs12328841 | 1.05E-08 | -0.03 | -0.03 - -0.02 | 0.422 |
| CDR | 9:13173885_G | MPDZ | rs4740546 | 1.06E-08 | 0.11 | 0.07 - 0.15 | 0.014 |
| CDR | 9:13181407_A | MPDZ | rs9696052 | 2.22E-08 | 0.11 | 0.07 - 0.14 | 0.015 |
| CDR | 9:13183256_A | MPDZ | rs10733255 | 3.18E-08 | 0.11 | 0.07 - 0.15 | 0.014 |
| CDR | 9:13194629_T | MPDZ | rs1831533 | 1.63E-08 | 0.11 | 0.07 - 0.15 | 0.013 |
| IOP_Baseline | 1:172221650_A | DNM3 | rs140007960 | 3.35E-08 | -1.14 | -1.54 - -0.73 | 0.025 |
| IOP_Baseline | 15:66836156_T | ZWILCH | rs116759068 | 1.72E-09 | -1.54 | -2.04 - -1.04 | 0.01 |
| IOP_Baseline | 18:56829888_G | LOC100286992 | rs151309448 | 3.12E-08 | -1.51 | -2.04 - -0.97 | 0.01 |
| IOP_Baseline | 1:93166853_G | EVI5 | rs115010249 | 4.70E-08 | -1.29 | -1.75 - -0.83 | 0.013 |
| IOP_Baseline | 2:42130567_A | LOC388942 | rs75115060 | 4.95E-10 | -1.92 | -2.52 - -1.31 | 0.01 |
| IOP_Baseline | 3:21924520_C | LOC100505836 | rs9816047 | 2.57E-08 | -1.19 | -1.61 - -0.77 | 0.021 |
| IOP_Baseline | 5:164031974_G | RPS15P6 | rs79763900 | 3.12E-10 | -1.58 | -2.07 - -1.09 | 0.011 |
| IOP_Baseline | 5:164044038_G | RPS15P6 | rs562005471 | 4.06E-10 | -1.56 | -2.05 - -1.07 | 0.011 |

| | | | | | | | |
|---|---|---|---|---|---|---|---|
| IOP_Baseline | 5:164063698_G | RPS15P6 | rs114606396 | 1.47E-09 | -1.49 | -1.98 - -1.01 | 0.012 |
| IOP_Baseline | 5:164066024_C | RPS15P6 | rs79796316 | 8.58E-09 | -1.44 | -1.93 - -0.95 | 0.012 |
| IOP_Baseline | 5:164109373_G | RPS15P6 | rs116200344 | 8.58E-09 | -1.44 | -1.93 - -0.95 | 0.012 |
| IOP_Baseline | 5:56080836_ATTATTTAT | MAP3K1 | | 5.13E-09 | -1.44 | -1.93 - -0.96 | 0.013 |
| IOP_Baseline | 6:18165033_A | TPMT | rs4716225 | 2.10E-08 | -1.2 | -1.62 - -0.78 | 0.021 |
| IOP_Baseline | 7:31217238_G | ADCYAP1R1 | rs150923992 | 3.08E-08 | -1.39 | -1.89 - -0.9 | 0.015 |
| IOP_Baseline | 7:31228984_A | ADCYAP1R1 | rs116683385 | 4.47E-08 | -1.34 | -1.82 - -0.86 | 0.016 |
| IOP_Baseline | 7:31239877_G | ADCYAP1R1 | rs137939838 | 2.99E-08 | -1.41 | -1.91 - -0.91 | 0.014 |
| IOP_Baseline | 7:31271691_C | NEUROD6 | rs115441482 | 1.64E-09 | -1.25 | -1.65 - -0.84 | 0.025 |
| IOP_Baseline | 7:31271964_T | NEUROD6 | rs143593389 | 2.20E-08 | -1.33 | -1.79 - -0.86 | 0.016 |
| IOP_Baseline | 7:97580066_A | OR7E7P | rs35195462 | 4.73E-08 | -0.74 | -1 - -0.47 | 0.068 |
| IOP_Baseline | 9:101815916_A | COL15A1 | rs114377702 | 1.11E-08 | -1.47 | -1.98 - -0.97 | 0.011 |
| IOP_Baseline | 9:101816103_A | COL15A1 | rs116365511 | 1.11E-08 | -1.47 | -1.98 - -0.97 | 0.011 |
| IOP_Max | 1:168291244_TAAG | TBX19 | | 1.05E-08 | -1.23 | -1.65 - -0.81 | 0.017 |
| IOP_Max | 1:172221650_A | DNM3 | rs140007960 | 4.19E-08 | -1.17 | -1.59 - -0.75 | 0.025 |
| IOP_Max | 13:114802008_T | RASA3 | rs568133975 | 1.55E-08 | -1.81 | -2.44 - -1.18 | 0.01 |
| IOP_Max | 14:33004728_C | AKAP6 | rs1956998 | 2.84E-08 | -1.28 | -1.73 - -0.83 | 0.017 |
| IOP_Max | 14:33008684_C | AKAP6 | rs76798482 | 1.93E-08 | -1.3 | -1.75 - -0.84 | 0.017 |
| IOP_Max | 15:66836156_T | ZWILCH | rs1167590683 | 4.20E-08 | -1.38 | -1.88 - -0.89 | 0.01 |
| IOP_Max | 16:78997237_G | WWOX | rs13336465 | 3.11E-08 | -1.24 | -1.68 - -0.8 | 0.016 |
| IOP_Max | 18:3112702_T | MYOM1 | rs142398219 | 4.77E-08 | -1.1 | -1.5 - -0.71 | 0.029 |
| IOP_Max | 18:56829888_G | LOC100286992 | rs151309448 | 1.72E-08 | -1.45 | -1.95 - -0.94 | 0.01 |
| IOP_Max | 19:47562980_T | ZC3H4 | rs116421545 | 2.02E-08 | -1.21 | -1.64 - -0.79 | 0.016 |
| IOP_Max | 2:218694006_C | TNS1 | rs112906382 | 1.67E-08 | -1.29 | -1.74 - -0.84 | 0.018 |
| IOP_Max | 2:42130567_A | LOC388942 | rs75115060 | 6.90E-10 | -1.84 | -2.42 - -1.25 | 0.01 |
| IOP_Max | 4:159470395_T | RXFP1 | rs573798101 | 7.54E-09 | -1.67 | -2.23 - -1.1 | 0.012 |
| IOP_Max | 4:159506013_T | RXFP1 | rs116536861 | 9.60E-09 | -1.67 | -2.24 - -1.1 | 0.012 |
| IOP_Max | 5:164031974_G | RPS15P6 | rs79763900 | 4.13E-11 | -1.6 | -2.07 - -1.12 | 0.011 |
| IOP_Max | 5:164044038_G | RPS15P6 | rs562005471 | 6.19E-09 | -1.48 | -1.99 - -0.98 | 0.011 |
| IOP_Max | 5:164063698_G | RPS15P6 | rs114606396 | 2.92E-08 | -1.39 | -1.88 - -0.9 | 0.012 |
| IOP_Max | 5:56080836_ATTATTTAT | MAP3K1 | | 1.72E-08 | -1.49 | -2.01 - -0.97 | 0.013 |
| IOP_Max | 6:18150250_T | TPMT | | 3.88E-09 | -1.19 | -1.59 - -0.79 | 0.023 |
| IOP_Max | 6:18165033_A | TPMT | rs47162255 | 3.25E-10 | -1.36 | -1.79 - -0.94 | 0.021 |
| IOP_Max | 6:18172270_C | KDM1B | rs16880324 | 1.56E-08 | -1.38 | -1.85 - -0.9 | 0.017 |
| IOP_Max | 6:20077425_G | MBOAT1 | rs112052428 | 2.36E-09 | -1.43 | -1.9 - -0.96 | 0.012 |
| IOP_Max | 6:76828118_T | IMPG1 | rs189745733 | 2.22E-08 | -1.08 | -1.46 - -0.7 | 0.029 |
| IOP_Max | 7:31213476_G | ADCYAP1R1 | rs146817296 | 8.07E-09 | -1.31 | -1.76 - -0.87 | 0.018 |
| IOP_Max | 7:31215363_AC | ADCYAP1R1 | | 2.32E-08 | -1.38 | -1.87 - -0.9 | 0.015 |
| IOP_Max | 7:31217238_G | ADCYAP1R1 | rs150923992 | 4.22E-09 | -1.45 | -1.93 - -0.96 | 0.015 |
| IOP_Max | 7:31228984_A | ADCYAP1R1 | rs116683385 | 3.15E-09 | -1.42 | -1.89 - -0.95 | 0.016 |
| IOP_Max | 7:31239877_G | ADCYAP1R1 | rs137939838 | 2.47E-09 | -1.51 | -2.01 - -1.02 | 0.014 |

| | | | | | | | |
|---|---|---|---|---|---|---|---|
| IOP_Max | 7:31271691_C | NEUROD6 | rs115441482 | 2.77E-10 | -1.35 | -1.77 - -0.93 | 0.025 |
| IOP_Max | 7:31271964_T | NEUROD6 | rs143593389 | 2.32E-09 | -1.43 | -1.9 - -0.96 | 0.016 |
| IOP_Max | 7:6066450_C | AIMP2 | rs2640 | 2.25E-08 | -1.29 | -1.74 - -0.84 | 0.018 |
| IOP_Max | 7:6079716_A | EIF2AK1 | rs74199305 | 2.80E-08 | -1.28 | -1.73 - -0.83 | 0.019 |
| IOP_Max | 7:6089026_G | EIF2AK1 | rs78429216 | 2.97E-08 | -1.21 | -1.64 - -0.78 | 0.022 |
| IOP_Max | 7:6109233_A | EIF2AK1 | rs79388869 | 2.64E-08 | -1.29 | -1.75 - -0.84 | 0.018 |
| IOP_Max | 9:101815916_A | COL15A1 | rs1143777026 | 1.21E-08 | -1.43 | -1.93 - -0.94 | 0.011 |
| IOP_Max | 9:101816103_A | COL15A1 | rs1163655116 | 1.21E-08 | -1.43 | -1.93 - -0.94 | 0.011 |
| MD | 10:12697942_T | MIR4481 | rs114483923 | 1.34E-09 | 3.63 | 2.46 - 4.81 | 0.012 |
| MD | 10:134625887_T | TTC40 | rs112986386 | 7.99E-09 | 3.39 | 2.24 - 4.54 | 0.017 |
| MD | 11:100832279_G | ARHGAP42 | rs150662590 | 1.68E-09 | 3.37 | 2.27 - 4.47 | 0.014 |
| MD | 11:77190251_C | PAK1 | rs145707826 | 1.62E-08 | 3.82 | 2.49 - 5.15 | 0.013 |
| MD | 11:77190646_G | DKFZp434E1119 | rs370650303 | 1.62E-08 | 3.82 | 2.49 - 5.15 | 0.013 |
| MD | 11:86721792_A | LOC100506368 | rs75095304 | 2.21E-08 | 3.28 | 2.13 - 4.44 | 0.011 |
| MD | 11:86793104_T | TMEM135 | rs77675473 | 1.31E-08 | 3.34 | 2.19 - 4.49 | 0.011 |
| MD | 11:98898299_T | CNTN5 | rs116643630 | 1.07E-08 | 3.13 | 2.05 - 4.2 | 0.02 |
| MD | 1:203918104_C | CBX1P3 | rs35108891 | 4.97E-08 | 3.31 | 2.12 - 4.5 | 0.011 |
| MD | 1:203920220_T | CBX1P3 | rs35473172 | 2.27E-08 | 3.11 | 2.02 - 4.2 | 0.016 |
| MD | 1:208046502_C | C1orf132 | rs80106999 | 3.88E-08 | 3.9 | 2.51 - 5.3 | 0.012 |
| MD | 12:105770611_C | C12orf75 | | 1.45E-08 | 3.02 | 1.98 - 4.07 | 0.014 |
| MD | 12:105774246_T | C12orf75 | | 1.45E-08 | 3.02 | 1.98 - 4.07 | 0.014 |
| MD | 1:217102298_A | ESRRG | rs114561905 | 3.95E-08 | 4.23 | 2.72 - 5.74 | 0.014 |
| MD | 1:238801736_G | MIPEPP2 | rs145925699 | 1.62E-08 | 2.98 | 1.94 - 4.01 | 0.02 |
| MD | 1:245130917_A | EFCAB2 | | 6.81E-10 | 4.17 | 2.85 - 5.5 | 0.011 |
| MD | 12:9944953_T | LOC100419929 | rs750461342 | 3.87E-08 | 3.25 | 2.09 - 4.41 | 0.023 |
| MD | 13:22575528_G | NME1P1 | rs142195625 | 2.24E-08 | 3.21 | 2.08 - 4.33 | 0.013 |
| MD | 13:26929723_CG | CDK8 | | 1.26E-08 | 3.07 | 2.01 - 4.13 | 0.02 |
| MD | 13:26965291_T | CDK8 | rs564628547 | 1.68E-08 | 3.04 | 1.98 - 4.1 | 0.02 |
| MD | 14:101687105_A | LOC100128373 | rs150298980 | 4.05E-09 | 3.99 | 2.66 - 5.32 | 0.012 |
| MD | 14:101706716_G | LOC100128373 | rs189996446 | 6.41E-10 | 3.77 | 2.57 - 4.97 | 0.011 |
| MD | 14:101753993_C | LOC100128373 | rs114831421 | 1.54E-08 | 3.62 | 2.37 - 4.88 | 0.013 |
| MD | 14:101755598_C | LOC100128373 | rs75726910 | 1.87E-08 | 3.74 | 2.44 - 5.05 | 0.011 |
| MD | 14:101762841_A | LOC100128373 | rs113888761 | 3.83E-09 | 3.84 | 2.56 - 5.12 | 0.013 |
| MD | 14:101771935_G | LOC100128373 | rs148365076 | 8.09E-10 | 4.03 | 2.74 - 5.32 | 0.013 |
| MD | 14:101773849_T | | | 4.96E-09 | 4 | 2.66 - 5.34 | 0.012 |
| MD | 14:101774096_A | LOC100128373 | rs149041531 | 6.14E-10 | 3.99 | 2.73 - 5.26 | 0.013 |
| MD | 14:101777003_A | LOC100128373 | rs149515116 | 1.32E-14 | 3.74 | 2.79 - 4.7 | 0.025 |
| MD | 14:41034514_G | LOC100533628 | rs142201506 | 9.48E-16 | 4.52 | 3.42 - 5.63 | 0.01 |
| MD | 14:41165668_G | LOC100533628 | rs75864954 | 6.83E-09 | 3.71 | 2.46 - 4.97 | 0.011 |
| MD | 14:45215389_G | DOCK11P1 | rs1091964 | 2.48E-09 | 3.99 | 2.68 - 5.31 | 0.012 |
| MD | 1:46731012_T | RAD54L | rs72901049 | 2.91E-11 | 3.57 | 2.52 - 4.62 | 0.015 |

| | | | | | | | |
|---|---|---|---|---|---|---|---|
| MD | 1:52402448_T | RAB3B | rs139624821 | 9.85E-09 | 3.92 | 2.58 - 5.26 | 0.013 |
| MD | 15:24531236_G | PWRN2 | rs35358454 | 3.97E-08 | 2.42 | 1.56 - 3.29 | 0.044 |
| MD | 15:35772993_G | ATPBD4 | rs185028035 | 2.58E-09 | 3.58 | 2.4 - 4.76 | 0.015 |
| MD | 15:35774389_A | ATPBD4 | rs114179978 | 2.58E-09 | 3.58 | 2.4 - 4.76 | 0.015 |
| MD | 15:46725050_G | LOC729316 | rs115008055 | 2.53E-09 | 3.98 | 2.67 - 5.29 | 0.014 |
| MD | 15:60559548_CAGA | ANXA2 | rs199994164 | 2.09E-08 | 3.28 | 2.13 - 4.43 | 0.019 |
| MD | 15:93822664_A | UNQ9370 | rs145053330 | 1.40E-11 | 4.09 | 2.9 - 5.28 | 0.014 |
| MD | 16:6388707_C | RBFOX1 | rs17139895 | 3.89E-08 | 3.79 | 2.43 - 5.14 | 0.011 |
| MD | 16:78343190_G | WWOX | rs80204300 | 3.09E-08 | 3.65 | 2.36 - 4.95 | 0.015 |
| MD | 1:76687603_T | ST6GALNAC3 | rs115818213 | 3.61E-11 | 3.74 | 2.63 - 4.85 | 0.01 |
| MD | 18:32988919_A | RPL7AP67 | rs181680212 | 6.12E-09 | 2.9 | 1.92 - 3.88 | 0.02 |
| MD | 18:33062491_C | INO80C | rs181851622 | 2.87E-09 | 3.03 | 2.03 - 4.03 | 0.019 |
| MD | 18:33067480_T | INO80C | | 1.43E-10 | 3.18 | 2.21 - 4.15 | 0.021 |
| MD | 18:33093734_A | INO80C | rs145594858 | 2.15E-09 | 3.06 | 2.06 - 4.06 | 0.02 |
| MD | 18:45851603_T | C18orf12 | rs116264801 | 1.27E-08 | 2.67 | 1.75 - 3.59 | 0.04 |
| MD | 18:45851792_A | C18orf12 | rs147641147 | 1.27E-08 | 2.67 | 1.75 - 3.59 | 0.04 |
| MD | 18:45851964_C | C18orf12 | rs114761171 | 3.42E-08 | 2.76 | 1.78 - 3.74 | 0.036 |
| MD | 18:45853761_T | C18orf12 | rs9960070 | 2.34E-08 | 2.78 | 1.8 - 3.75 | 0.036 |
| MD | 18:45856557_G | C18orf12 | rs144488026 | 2.39E-08 | 2.88 | 1.87 - 3.89 | 0.035 |
| MD | 18:75616220_T | LOC100421527 | rs117283685 | 1.17E-08 | 3.37 | 2.21 - 4.53 | 0.012 |
| MD | 18:75616859_T | LOC100421527 | rs117460825 | 1.17E-08 | 3.37 | 2.21 - 4.53 | 0.012 |
| MD | 18:75622172_A | LOC100421527 | rs4890863 | 6.69E-09 | 3.35 | 2.22 - 4.49 | 0.01 |
| MD | 18:75697997_C | LOC100421527 | | 8.51E-12 | 4.01 | 2.86 - 5.17 | 0.01 |
| MD | 19:22610632_T | ZNF98 | rs535930483 | 1.95E-09 | 4.33 | 2.91 - 5.74 | 0.013 |
| MD | 19:22616386_C | ZNF98 | rs188375694 | 1.95E-09 | 4.33 | 2.91 - 5.74 | 0.013 |
| MD | 19:22625703_T | ZNF98 | rs191540610 | 3.12E-08 | 4.28 | 2.77 - 5.8 | 0.012 |
| MD | 19:22627656_G | ZNF209P | rs151038343 | 3.12E-08 | 4.28 | 2.77 - 5.8 | 0.012 |
| MD | 19:22636874_A | ZNF209P | rs143227593 | 3.12E-08 | 4.28 | 2.77 - 5.8 | 0.012 |
| MD | 19:22639568_T | ZNF209P | rs143123340 | 3.12E-08 | 4.28 | 2.77 - 5.8 | 0.012 |
| MD | 19:34233029_G | CHST8 | rs71351737 | 4.51E-09 | 3.7 | 2.46 - 4.93 | 0.012 |
| MD | 20:14353817_A | MACROD2 | rs370794768 | 2.34E-08 | 3.6 | 2.33 - 4.86 | 0.013 |
| MD | 20:14358134_G | MACROD2 | rs76211883 | 2.34E-08 | 3.6 | 2.33 - 4.86 | 0.013 |
| MD | 20:14362754_A | MACROD2 | rs113662958 | 2.34E-08 | 3.6 | 2.33 - 4.86 | 0.013 |
| MD | 20:14376543_G | MACROD2 | rs113188443 | 7.22E-09 | 3.8 | 2.51 - 5.08 | 0.012 |
| MD | 20:15666084_G | MACROD2 | rs6110735 | 6.95E-09 | 2.91 | 1.93 - 3.9 | 0.03 |
| MD | 20:15674785_T | MACROD2 | rs117074103 | 5.65E-10 | 3.11 | 2.13 - 4.1 | 0.03 |
| MD | 20:15916938_A | MACROD2 | rs181472575 | 3.27E-08 | 3.52 | 2.27 - 4.76 | 0.012 |
| MD | 20:58191696_A | PHACTR3 | rs141640824 | 2.13E-11 | 4.35 | 3.07 - 5.62 | 0.01 |
| MD | 2:109843921_C | SH3RF3 | rs17269668 | 3.30E-08 | 3.71 | 2.39 - 5.03 | 0.012 |
| MD | 21:18172016_G | LINC00478 | rs191063909 | 4.95E-08 | 3.56 | 2.28 - 4.83 | 0.017 |
| MD | 21:18173803_A | LINC00478 | rs112551494 | 4.95E-08 | 3.56 | 2.28 - 4.83 | 0.017 |

| | | | | | | | |
|---|---|---|---|---|---|---|---|
| MD | 21:18175117_G | LINC00478 | rs113502263 | 4.95E-08 | 3.56 | 2.28 - 4.83 | 0.017 |
| MD | 21:18176034_A | LINC00478 | rs113637243 | 4.95E-08 | 3.56 | 2.28 - 4.83 | 0.017 |
| MD | 21:18176826_T | LINC00478 | rs113123910 | 1.72E-08 | 3.71 | 2.42 - 5 | 0.017 |
| MD | 21:18178771_T | LINC00478 | rs74424479 | 4.95E-08 | 3.56 | 2.28 - 4.83 | 0.017 |
| MD | 21:18196762_A | LINC00478 | rs113557325 | 3.63E-08 | 3.61 | 2.33 - 4.9 | 0.017 |
| MD | 21:18198836_A | LINC00478 | rs148605754 | 3.63E-08 | 3.61 | 2.33 - 4.9 | 0.017 |
| MD | 21:18199603_A | LINC00478 | rs113028845 | 2.25E-08 | 3.64 | 2.37 - 4.92 | 0.017 |
| MD | 21:18202647_A | LINC00478 | rs150145974 | 3.63E-08 | 3.61 | 2.33 - 4.9 | 0.017 |
| MD | 2:212236240_T | ERBB4 | rs77593502 | 1.06E-09 | 3.62 | 2.45 - 4.78 | 0.012 |
| MD | 2:212273739_T | ERBB4 | rs112122313 | 6.59E-10 | 3.7 | 2.52 - 4.87 | 0.012 |
| MD | 2:232200749_T | ARMC9 | rs146931289 | 2.50E-09 | 4 | 2.69 - 5.32 | 0.014 |
| MD | 22:50321429_A | ALG12 | rs75995685 | 2.15E-10 | 3.92 | 2.71 - 5.13 | 0.014 |
| MD | 3:113281021_C | SIDT1 | rs143713583 | 9.78E-09 | 3.85 | 2.54 - 5.17 | 0.012 |
| MD | 3:136863770_A | IL20RB | | 2.23E-08 | 2.81 | 1.82 - 3.79 | 0.025 |
| MD | 3:142293247_C | ATR | rs116088897 | 6.59E-09 | 2.78 | 1.84 - 3.72 | 0.024 |
| MD | 4:131963445_T | GAPDHP56 | rs17051142 | 7.16E-10 | 3.78 | 2.58 - 4.99 | 0.014 |
| MD | 4:132016166_AT | GAPDHP56 | | 9.02E-09 | 3.48 | 2.29 - 4.66 | 0.017 |
| MD | 4:137802259_A | SERF1AP1 | rs145330103 | 3.33E-08 | 3.4 | 2.19 - 4.6 | 0.013 |
| MD | 4:27213164_G | STIM2 | rs116570872 | 2.41E-08 | 3.68 | 2.39 - 4.98 | 0.013 |
| MD | 4:27214236_C | STIM2 | rs75485894 | 2.41E-08 | 3.68 | 2.39 - 4.98 | 0.013 |
| MD | 5:112821871_G | MCC | rs111680425 | 2.96E-08 | 2.98 | 1.92 - 4.03 | 0.03 |
| MD | 5:41563196_T | TCP1P2 | rs188486862 | 3.94E-08 | 3.5 | 2.25 - 4.75 | 0.011 |
| MD | 5:41563987_G | TCP1P2 | rs148598417 | 3.94E-08 | 3.5 | 2.25 - 4.75 | 0.011 |
| MD | 5:41577596_G | TCP1P2 | rs143179640 | 3.94E-08 | 3.5 | 2.25 - 4.75 | 0.011 |
| MD | 5:42093354_G | LOC100130042 | rs146433882 | 1.09E-09 | 3.97 | 2.7 - 5.25 | 0.011 |
| MD | 6:113881248_A | RPS27AP11 | rs9488212 | 1.50E-08 | 2.56 | 1.67 - 3.45 | 0.045 |
| MD | 6:114602648_C | LOC441167 | rs185376250 | 4.73E-08 | 3.38 | 2.16 - 4.59 | 0.01 |
| MD | 6:23341967_A | RPL6P18 | rs144256194 | 4.73E-09 | 3.33 | 2.21 - 4.44 | 0.019 |
| MD | 6:70670418_C | COL19A1 | rs115364550 | 3.76E-08 | 3.75 | 2.41 - 5.08 | 0.011 |
| MD | 6:96018150_A | MANEA | rs116680063 | 1.91E-08 | 2.83 | 1.84 - 3.82 | 0.021 |
| MD | 6:96087322_AC | MANEA | | 1.64E-08 | 2.63 | 1.72 - 3.55 | 0.028 |
| MD | 7:11542978_G | THSD7A | rs6980010 | 3.36E-08 | 3.32 | 2.14 - 4.5 | 0.026 |
| MD | 7:13215091_G | RBMX2P4 | rs148116853 | 1.13E-08 | 3.71 | 2.44 - 4.98 | 0.011 |
| MD | 7:13239462_G | RBMX2P4 | rs74773569 | 1.64E-09 | 3.63 | 2.45 - 4.81 | 0.011 |
| MD | 7:13241140_C | RBMX2P4 | rs143620602 | 1.64E-09 | 3.63 | 2.45 - 4.81 | 0.011 |
| MD | 7:25368343_T | NPVF | rs112700836 | 1.96E-08 | 3.23 | 2.1 - 4.35 | 0.012 |
| MD | 7:25369213_A | NPVF | rs73283010 | 1.96E-08 | 3.23 | 2.1 - 4.35 | 0.011 |
| MD | 7:25369823_A | NPVF | rs73283011 | 1.96E-08 | 3.23 | 2.1 - 4.35 | 0.012 |
| MD | 7:25370548_C | NPVF | rs73283013 | 1.96E-08 | 3.23 | 2.1 - 4.35 | 0.012 |
| MD | 7:27882316_T | JAZF1 | rs62451072 | 2.88E-08 | 3.51 | 2.27 - 4.76 | 0.013 |
| MD | 7:27882471_C | JAZF1 | rs10486562 | 2.88E-08 | 3.51 | 2.27 - 4.76 | 0.013 |

| | | | | | | | |
|---|---|---|---|---|---|---|---|
| MD | 7:5266629_C | WIPI2 | rs59864675 | 2.42E-09 | 3.37 | 2.26 - 4.48 | 0.016 |
| MD | 8:25565605_A | LOC100507222 | rs114396168 | 1.77E-08 | 3.1 | 2.02 - 4.17 | 0.018 |
| MD | 8:53938061_T | LOC101060191 | rs116452158 | 3.25E-13 | 4.21 | 3.07 - 5.34 | 0.011 |
| MD | 8:75686918_C | FLJ39080 | rs545831765 | 3.91E-09 | 4.18 | 2.79 - 5.57 | 0.011 |
| MD | 8:75699089_T | FLJ39080 | rs192962111 | 4.68E-10 | 4.3 | 2.94 - 5.65 | 0.012 |
| MD | 8:77301021_A | MRPL9P1 | rs16939276 | 3.87E-09 | 3.82 | 2.55 - 5.1 | 0.01 |
| MD | 9:129436105_A | LMX1B | rs143217136 | 1.12E-08 | 3.07 | 2.02 - 4.13 | 0.012 |
| MD | 9:129439050_T | LMX1B | rs115683895 | 7.42E-09 | 3.06 | 2.02 - 4.1 | 0.012 |
| MD | 9:129446114_A | LMX1B | rs144229999 | 2.24E-10 | 3.23 | 2.23 - 4.22 | 0.01 |
| MD | 9:129446648_C | LMX1B | rs147720587 | 5.28E-13 | 3.51 | 2.56 - 4.47 | 0.012 |
| MD | 9:129448920_T | LMX1B | rs140140891 | 5.28E-13 | 3.51 | 2.56 - 4.47 | 0.012 |
| MD | 9:129449650_C | LMX1B | rs187699205 | 5.28E-13 | 3.51 | 2.56 - 4.47 | 0.012 |
| MD | 9:91279580_A | LOC286238 | | 8.54E-09 | 3.6 | 2.37 - 4.83 | 0.01 |
| MD | 9:91284380_T | LOC286238 | rs181197053 | 2.92E-08 | 3.54 | 2.29 - 4.79 | 0.01 |
| PSD | 10:113517524_A | RPS6P15 | rs142087346 | 1.78E-09 | -1.38 | -1.83 - -0.93 | 0.015 |
| PSD | 10:113528369_C | RPS6P15 | rs145960357 | 1.08E-08 | -1.33 | -1.79 - -0.88 | 0.016 |
| PSD | 10:113529507_A | RPS6P15 | rs138116457 | 7.35E-09 | -1.36 | -1.83 - -0.9 | 0.015 |
| PSD | 10:113535805_C | RPS6P15 | rs114064009 | 1.79E-08 | -1.34 | -1.8 - -0.87 | 0.015 |
| PSD | 10:99390194_C | MORN4 | rs111941669 | 1.44E-09 | -1.52 | -2.01 - -1.02 | 0.016 |
| PSD | 10:99395853_A | MORN4 | rs113671174 | 3.74E-09 | -1.49 | -1.98 - -0.99 | 0.015 |
| PSD | 10:99397796_T | MORN4 | rs111997721 | 1.51E-08 | -1.47 | -1.97 - -0.96 | 0.015 |
| PSD | 11:77190251_C | DKFZp434E111 9 | rs145707826 | 3.48E-10 | -1.44 | -1.89 - -0.99 | 0.013 |
| PSD | 11:77190646_G | PAK1 | rs370650303 | 3.48E-10 | -1.44 | -1.89 - -0.99 | 0.013 |
| PSD | 11:8385488_A | STK33 | rs151244129 | 4.76E-08 | -1.5 | -2.04 - -0.96 | 0.011 |
| PSD | 11:8392761_C | STK33 | rs146828273 | 2.11E-09 | -1.64 | -2.18 - -1.11 | 0.012 |
| PSD | 11:8394373_A | STK33 | rs533094261 | 8.30E-11 | -1.75 | -2.28 - -1.22 | 0.012 |
| PSD | 11:8401590_A | STK33 | rs190526296 | 2.28E-10 | -1.77 | -2.31 - -1.22 | 0.01 |
| PSD | 11:8401675_T | STK33 | rs569329344 | 2.28E-10 | -1.77 | -2.31 - -1.22 | 0.01 |
| PSD | 11:8418450_T | STK33 | rs569683432 | 3.20E-09 | -1.69 | -2.24 - -1.13 | 0.01 |
| PSD | 11:8419845_A | STK33 | rs553348782 | 5.29E-09 | -1.65 | -2.2 - -1.09 | 0.01 |
| PSD | 11:8421097_G | STK33 | rs534038252 | 5.29E-09 | -1.65 | -2.2 - -1.09 | 0.01 |
| PSD | 14:21657929_A | LOC100129923 | rs182556456 | 3.08E-08 | -1.61 | -2.18 - -1.04 | 0.011 |
| PSD | 17:1425146_A | PITPNA | rs75951666 | 1.39E-09 | -1.65 | -2.19 - -1.12 | 0.011 |
| PSD | 17:28566663_T | BLMH | rs116657864 | 2.11E-11 | -1.61 | -2.08 - -1.14 | 0.01 |
| PSD | 17:28600117_GT | BLMH | | 2.11E-11 | -1.61 | -2.08 - -1.14 | 0.011 |
| PSD | 17:28602192_G | BLMH | rs139315505 | 9.62E-12 | -1.62 | -2.08 - -1.15 | 0.011 |
| PSD | 17:39248459_T | KRTAP4-7 | rs148001047 | 1.02E-13 | -2.09 | -2.64 - -1.54 | 0.01 |
| PSD | 17:72848018_T | GRIN2C | rs148188891 | 4.06E-09 | -1.48 | -1.97 - -0.99 | 0.016 |
| PSD | 17:72856560_T | FDXR | rs114802931 | 8.39E-09 | -1.44 | -1.93 - -0.95 | 0.017 |
| PSD | 20:48841343_G | CEBPB | rs572657086 | 3.89E-09 | -1.48 | -1.98 - -0.99 | 0.011 |
| PSD | 21:18879171_T | RPL39P40 | rs139461536 | 3.49E-08 | -1.32 | -1.79 - -0.85 | 0.015 |

| | | | | | | | |
|---|---|---|---|---|---|---|---|
| PSD | 2:148342366_C | RPL26P14 | rs116575189 | 7.94E-10 | -1.11 | -1.46 - -0.75 | 0.03 |
| PSD | 2:15127499_A | NBAS | rs189692504 | 3.98E-11 | -1.54 | -2 - -1.08 | 0.011 |
| PSD | 4:126984908_T | LOC100419353 | rs74624148 | 1.34E-08 | -1.56 | -2.1 - -1.02 | 0.011 |
| PSD | 4:136533571_A | TARS2P1 | rs114437581 | 4.88E-08 | -0.89 | -1.21 - -0.57 | 0.052 |
| PSD | 4:26216024_AAGAG | LOC645481 | | 4.37E-08 | -1.3 | -1.77 - -0.84 | 0.018 |
| PSD | 5:107160240_A | FBXL17 | rs79012700 | 4.80E-08 | -1.16 | -1.57 - -0.74 | 0.031 |
| PSD | 5:53262321_G | ARL15 | rs6879379 | 7.00E-10 | -1.28 | -1.69 - -0.87 | 0.02 |
| PSD | 5:53270084_C | ARL15 | | 3.93E-08 | -1.19 | -1.62 - -0.77 | 0.02 |
| PSD | 5:53271410_G | ARL15 | rs73754403 | 3.93E-08 | -1.19 | -1.62 - -0.77 | 0.02 |
| PSD | 6:170543043_G | LOC154449 | rs111409772 | 8.03E-09 | -1.52 | -2.03 - -1 | 0.014 |
| PSD | 6:170543059_A | LOC154449 | rs147948513 | 8.03E-09 | -1.52 | -2.03 - -1 | 0.014 |
| PSD | 6:37759865_T | ZFAND3 | | 6.00E-10 | -1.24 | -1.63 - -0.85 | 0.024 |
| PSD | 7:136236414_A | LOC100421642 | rs140122465 | 2.29E-08 | -1.67 | -2.26 - -1.08 | 0.011 |
| PSD | 7:136256121_T | LOC100421642 | rs115205488 | 4.00E-08 | -1.66 | -2.25 - -1.06 | 0.011 |
| PSD | 7:19796507_T | TMEM196 | rs79016864 | 2.12E-10 | -1.66 | -2.17 - -1.14 | 0.011 |
| PSD | 9:120198167_G | ASTN2 | | 4.18E-08 | -1.55 | -2.11 - -1 | 0.011 |
| RNFL | 2:109560289_A | EDAR | rs114101591 | 9.19E-09 | -8.36 | -11.22 - -5.51 | 0.014 |

*Mapped/nearest gene=variant in gene region, with nearest gene shaded
rsID: dbSNP ID, Beta=effect size, CI=confidence interval, CDR=cup-to-disc ratio, IOP=intraocular pressure, MD=mean deviation, PSD=pattern standard deviation, RNFL=retinal nerve fiber layer

#### Extended Data Table. 4:

**Table 4a:** Table showing expression of the genes in each locus that comprise over 95% of associated MAGMA gene variants in adult human eye tissues, as obtained from the Ocular Tissue Database (OTDB), along with gene expression values. In the OTDB, gene expression is represented as an Affymetrix Probe Logarithmic Intensity Error (PLIER) number for an individual probe. These numbers were calculated as described in Wagner *et al*., 2013.

| Gene | RPE Choroid | Ciliary Body | Cornea | Iris | Lens | Optic Nerve | Optic Nerve Head | Retina | Sclera | Trabecular Meshwork |
|---|---|---|---|---|---|---|---|---|---|---|
| *CLCF1* | 39.8 | 32.0 | 41.0 | 38.8 | 40.0 | 42.0 | 33.6 | 47.2 | 38.8 | 44.9 |
| *STXBP3* | 73.2 | 95.8 | 162.8 | 93.5 | 238.7 | 242.1 | 314.4 | 153.7 | 79.3 | 193.3 |
| *LAMB1* | 146.7 | 136.9 | 183.7 | 144.3 | 131.7 | 60.2 | 137.9 | 32.7 | 48.8 | 257.5 |
| *TLN2* | 31.9 | 31.3 | 38.3 | 27.3 | 53.0 | 58.0 | 54.9 | 47.3 | 34.7 | 28.8 |
| *DNAJC5B* | 19.5 | 26.0 | 23.1 | 23.4 | 17.7 | 17.8 | 20.5 | 17.3 | 21.4 | 26.9 |
| *CMTR1/ FTSJD2* | 24.6 | 27.6 | 40.9 | 19.7 | 39.4 | 37.5 | 48.6 | 43.4 | 34.3 | 53.6 |
| *FAM209A/C20orf106* | 72.5 | 76.4 | 95.0 | 96.2 | 138.5 | 110.0 | 105.5 | 66.1 | 93.3 | 100.7 |

**Table 4b:** Gene expression levels MAGMA gene variants in mouse ONH

| | | | STAGE 1 | | STAGE 2 | | STAGE 3 | | STAGE 4 | | STAGE 5 | |
|---|---|---|---|---|---|---|---|---|---|---|---|---|
| **Gene** | **Probe Set ID** | **Probe Sets#** | **FC** | **QV** | **FC** | **QV** | **FC** | **QV** | **FC** | **QV** | **FC** | **QV** |
| *Clcf1* | 1437270_a_at | 3 | 1.434 | 0.124 | 1.386 | 0.011 | **2.170** | **0.000** | **4.408** | **0.000** | 1.272 | 0.128 |
| *Stxbp3* | 1416653_at | 1 | 1.209 | 0.476 | 1.050 | 0.470 | 1.009 | 0.529 | 1.350 | 0.079 | 1.466 | 0.024 |
| *Lamb1* | 1424113_at | 4 | 1.070 | 0.699 | 1.367 | 0.017 | 1.273 | 0.049 | 1.624 | 0.017 | 1.100 | 0.341 |
| *Tln2* | 1435700_at | 3 | -1.100 | 0.489 | **-1.270** | **0.006** | **-1.304** | **0.005** | **-1.879** | **0.001** | **-2.010** | **0.001** |
| *Dnajc5b* | 1418725_at | 1 | 1.019 | 0.777 | 1.050 | 0.464 | 1.008 | 0.526 | 1.265 | 0.026 | 1.186 | 0.104 |
| *Fam209* | 1429544_at | 1 | -1.030 | 0.735 | 1.082 | 0.338 | 1.022 | 0.473 | 1.091 | 0.355 | 1.314 | 0.020 |

The above table shows previously published microarray gene expression dataset in optic nerve head punches and was obtained from submitting data into the OTDB database server. The Log2 fold change (FC) shown here compares DBA/2J mice relative to DBA/2J-Gpnmb+ controls at 5 molecularly distinct stages of glaucoma defined using hierarchical clustering. Stg- Stages 1, 2 and 3 are early states of glaucoma, Stages 4 and 5 have eyes with moderate and severe optic nerve damage respectively. Q-value (QV) shows the significance of the FC of GWAS genes at each stage. Bold font represents significant expression.
